# Hippo Signaling Cofactor, WWTR1, at the Crossroads of Human Trophoblast Progenitor Self-Renewal and Differentiation

**DOI:** 10.1101/2022.03.06.483207

**Authors:** Soma Ray, Abhik Saha, Ananya Ghosh, Namrata Roy, Ram P Kumar, Gudrun Meinhardt, Abhirup Mukerjee, Sumedha Gunewardena, Rajnish Kumar, Martin Knöfler, Soumen Paul

## Abstract

Healthy progression of human pregnancy relies on cytotrophoblast progenitor (CTB) self-renewal and their differentiation towards multi-nucleated syncytiotrophoblasts (STBs) and invasive extravillous trophoblasts (EVTs). However, underlying molecular mechanisms that fine-tune CTB self-renewal or direct their differentiation towards STBs or EVTs during human placentation are poorly defined. Here, we show that hippo signaling cofactor WW Domain Containing Transcription Regulator 1 (WWTR1) is a master regulator of trophoblast fate choice during human placentation. Using human trophoblast stem cells (human TSCs), primary CTBs and human placental explants, we demonstrate that WWTR1 promotes self-renewal in human CTBs and is essential for their differentiation to EVTs. In contrast, WWTR1 prevents induction of STB fate in undifferentiated CTBs. Our single-cell RNA-sequencing analyses in first-trimester human placenta along with mechanistic analyses in human TSCs revealed that WWTR1 fine-tunes trophoblast fate by directly regulating Wnt signaling components. Importantly, our analyses of placentae from pathological pregnancies show that extreme preterm birth (gestational time ≤28weeks) and intrauterine growth restriction along with preeclampsia (IUGR/PE) are often associated with loss of WWTR1 expression in CTBs. In summary, our findings establish a critical importance of WWTR1 at the crossroads of human trophoblast progenitor self-renewal vs. differentiation. It plays positive instructive roles to promote CTB self-renewal and EVT differentiation and safeguards undifferentiated CTBs from obtaining the STB fate.

**SIGNIFICANCE:** Human pregnancy relies on formation of the transient organ placenta and trophoblast cells are the major building blocks of the placenta. A defect in trophoblast progenitor self-renewal or their differentiation is associated with either pregnancy loss or pathological pregnancies, yet underlying molecular mechanisms that regulate trophoblast differentiation are poorly understood. In this study, we discovered that WWTR1, a transcription cofactor and a component of conserved Hippo signaling pathway, optimizes trophoblast progenitor self-renewal and is essential for their differentiation into the invasive extravillous trophoblast cell lineage. Our findings establish WWTR1 as a critical regulator for success in human placentation and progression of a healthy pregnancy.

## INTRODUCTION

Establishment of human pregnancy is associated with formation of an invasive primitive syncytium from CTB progenitors at the blastocyst implantation site (1-3). Subsequently, proliferation and differentiation of CTB progenitors result in formation of the functional villous placenta, containing two types of matured villi; (i) floating villi, which float in the maternal blood and contain the STB population that establish the maternal-fetal nutrient and gas exchange interface and secretes human chorionic gonadotropin to maintain corpus luteum (4, 5), and (ii) anchoring villi, which anchor to maternal tissue and contain the invasive EVT population (Fig. 1A). In anchoring villi, CTB progenitors adapt a distinct differentiation pathway. At the base of the anchoring villi, CTB progenitors proliferate to form a CTB cell column. Eventually, cells at the distal ends of the CTB column differentiate to adapt a migratory phenotype, thereby establishing the invasive EVT lineage, which orchestrates the uterine environment, expresses non-classical human lecukocyte antigen (HLA)-G and promotes immune tolerance to the fetus to secure progression of pregnancy (5-7). A subset of EVTs invades and remodels the uterine vasculature to establish enhanced maternal blood supply at the uterine-placental interface to fulfill the nutrient requirement of the growing fetus (8). Thus, self-renewal of CTB progenitors and their differentiation to STBs and EVTs in floating vs. anchoring villi are the essential events for progression of human pregnancy. Failures in these processes are implicated in early pregnancy loss or pregnancy-associated diseases such as preeclampsia, intrauterine growth restriction, and preterm birth (9-14).

**Figure 1:**
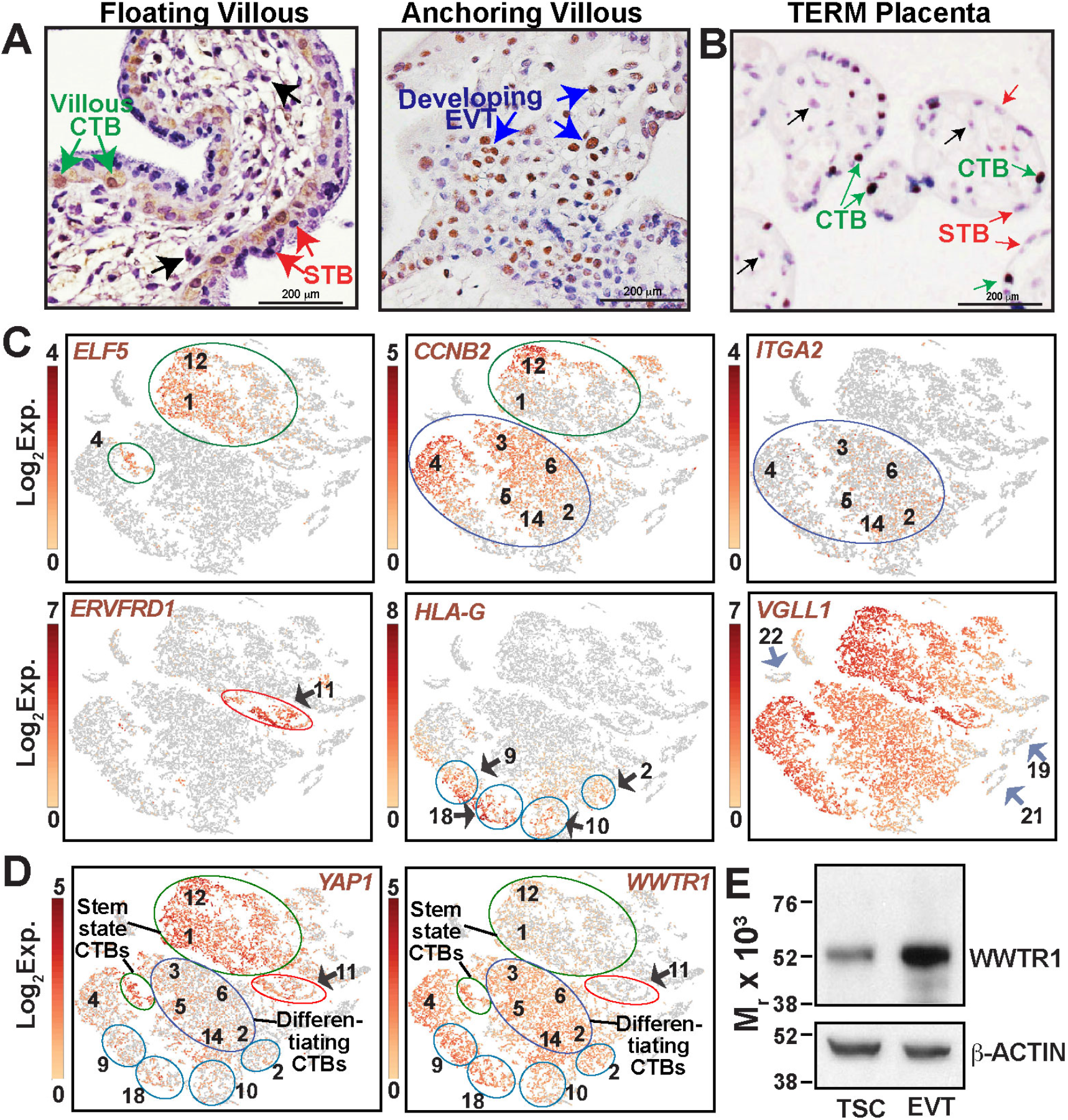
WWTR1 expression in human placentation site. (A) Immunostained images show WWTR1 protein expression in a first-trimester (week 8) human placental villous. WWTR1 expression was detected in CTBs within floating villi (green arrows) and in developing EVTs (blue arrows) in anchoring villi. However, WWTR1 protein expression was undetectable in STBs (red arrows) and in stromal cells (black arrows) within floating villi. (B) Immunostained image shows WWTR1 protein expression in a normal term human placenta. WWTR1 is expressed in CTBs (green arrows). STBs (red arrows) and stromal cells (black arrows) do not express WWTR1. (C) t-SNE plots of the single-cell clusters of first-trimester human placental samples. Expressions of specific genes were monitored to identify clusters of different CTB progenitors and developing EVTs. (D) t-SNE plots showing differential mRNA expression patterns of *YAP1* and *WWTR1* in single cell clusters, obtained by scRNA-seq analyses in first-trimester human placentae. YAP1 is highly expressed in clusters of stem-state CTBs (green circles) and in CTBs of cluster 11 (red circle), which are mitotically arrested. In contrast, WWTR1 expression induced in mitotically active but differentiating CTBs and in cells of cluster 9,10, 18, and a part of clusters 2 (blue circles), which represent developing EVTs. (E) Western blots show induction of WWTR1 protein expression in human TSCs upon differentiation to EVTs.

Although CTBs establish the stem/progenitor compartment of a developing placenta, recent studies revealed that distinct populations of CTBs exist within a first-trimester placenta. *Ex vivo* developmental analyses of peri-implantation human embryos and global gene expression analyses including single-cell mRNA sequencing (scRNA-seq) revealed that during early stages of human placentation, along with undifferentiated CTBs, populations of mitotically active CTBs arise that are poised for either STB or EVT differentiation (15-17). In floating villi of the early first-trimester placenta (4-8 weeks), expression of CDX2 and ELF5 marks the undifferentiated stem-state CTBs (18, 19), which are not committed to the differentiation pathway, whereas mitotically active but differentiating CTBs can be identified by the expression of genes that are linked to the interferon response, like interferon gamma receptor 2 (IFNGR2), and cell cycle regulators such as CDK1 and CCNB2 (15, 16). In addition, a population of mitotically inactive CTBs, which are committed for STB differentiation, can be identified by the expression of retroviral protein ERVFRD1 (16). Unlike in floating villi, CDX2 expression is suppressed in column CTBs within anchoring villi and ELF5 mRNA is expressed only in column CTBs near the base of the cell column (proximal column). Thus, it was proposed that ELF5 transcriptional activity regulates the trophoblast stem-state compartment of a developing human placenta (18). The undifferentiated column CTB subpopulation expresses integrin A2 (ITGA2) and NOTCH1 (20, 21). The transition of column CTBs to EVTs is associated with loss of ITGA2 expression and induction of specific genes, such as *ITGA1, MMP2* and human leukocyte antigen *HLA-G* (22-24). Thus, CTB self-renewal and differentiation during human placentation is a highly dynamic process and relies on molecular mechanisms that fine-tune the gene expression programs in different CTB progenitor subpopulations.

Molecular mechanisms that regulate trophoblast development during early human placentation were poorly understood due to ethical restrictions and lack of appropriate model systems. However, successful derivation of human trophoblast stem cells from CTBs of first-trimester human placentae (25) and success in CTB-organoid culture (26, 27) have provided excellent models to identify regulatory pathways that are involved in CTB self-renewal and their differentiation to STBs and EVTs. Using human TSCs and CTB organoids, we identified conserved Hippo signaling components, transcription factor TEAD4 and cofactor YAP1, as important regulators to maintain the self-renewal ability of CTBs in a developing human placenta (11, 28). We showed that TEAD4 and YAP1 are selectively expressed in undifferentiated CTBs and loss of either TEAD4 or YAP1 in CTBs impairs their self-renewal ability (11, 28).

Along with YAP1, TEAD4 can interact with other cofactors, such as VGLL1 and WWTR1, to regulate gene expression programs. Earlier study identified VGLL1 as a human-specific marker of proliferative CTBs and proposed a regulatory role for VGLL1 in the TEAD4-mediated gene expression program during human trophoblast lineage development (19). However, the importance of WWTR1, a paralog of YAP1 and VGLL1 and another major cofactor of the Hippo signaling pathway, in human trophoblast development is yet to be defined.

Interestingly, studies focusing on trophoblast lineage development in mouse and marsupials indicated that YAP1 and WWTR1 might have redundant or mutually distinct roles during trophoblast development. Studies with YAP1 and WWTR1 mutant mouse models revealed a redundant role in trophoblast lineage development. Although early trophoblast development was not impaired in either *Wwtr1^−/−^* or *Yap1^−/−^* embryos, the *Yap1/Wwtr1* double knockout embryos failed to form blastocysts due to defective development of the trophectoderm lineage (29). In contrast to mouse, trophoblast cells of a developing marsupial embryo show distinct expression patterns of YAP1 and WWTR1. In marsupials, WWTR1 expression is strongly maintained within the nuclei of developing trophoblast lineage, whereas YAP1 expression is suppressed. Thus, it was predicted that WWTR1, but not YAP1, has a more important role in trophoblast maintenance in a developing marsupial embryo (30). Together, these studies indicated that WWTR1, either in conjunction with YAP1 or in an independent fashion, contributes to the trophoblast lineage development during mammalian placentation.

Here, using human TSCs, primary CTBs and placental explants, we have focused our attention on the functional importance of WWTR1 in human trophoblast development, especially in the context of CTB self-renewal and their differentiation to STBs and EVTs. In addition, we examined possible correlation of defective WWTR1 function in pathological pregnancies. We discovered that, similar to YAP1, WWTR1 is required to maintain self-renewal of CTB progenitors. In addition, WWTR1 is important to prevent STB differentiation and to induce the EVT differentiation program in CTBs. We also found that pregnancies associated with extreme preterm birth as well as preterm birth along with IUGR/PE are often associated with loss of WWTR1 in CTBs. Collectively, our findings implicate WWTR1 as an important orchestrator of trophoblast development during human placentation.

## RESULTS

### During human placentation WWTR1 is expressed in CTB progenitors and the expression is induced during EVT development

To define the importance of WWTR1 in human trophoblast development, we tested WWTR1 protein expressions in first-trimester human placentae (6-8 weeks of gestation). As mentioned earlier, floating villi in a first-trimester human placenta contain two different layers of trophoblast cells: (i) a layer of CTB progenitors and (ii) the post-mitotic STB layer, overlaying the CTBs. In contrast, the anchoring villi contain the CTB column and cells at the distal end of the CTB columns develop to the invasive EVTs. We found that WWTR1 is predominantly expressed in CTBs within floating villi (Fig. 1A, green arrows). We also noticed WWTR1 expression in emerging EVTs within anchoring villi (Fig. 1A, blue arrows). However, WWTR1 expression is suppressed in differentiated STBs (Fig. 1A, red arrows) and in stromal cells (Fig. 1A, black arrows) within floating villi. We also tested WWTR1 expression in term human placentae. Similar to first-trimester floating villi, WWTR1 is only expressed in CTBs (Fig. 1B, green arrows) in a normal term human placenta and is repressed in STBs (Fig. 1B, red arrows) and in stromal cells (Fig. 1B, black arrows).

Single-cell RNA-sequencing (scRNA-seq) analyses with first-trimester human placentae revealed that a developing human placenta contains distinct CTB subpopulations, which could be identified via expression of specific genes (16). We hypothesized that WWTR1 and other Hippo signaling cofactors, namely YAP1 and VGLL1, might have distinct functions in different CTB subpopulations of a developing human placenta. Therefore, we compared expression of *WWTR1*, *YAP1* and *VGLL1* in different CTB subpopulations. To that end, we analyzed scRNA-seq data that we generated with first-trimester human placentae. Based on gene expression patterns, entire single cell populations of 6-8-week human placentae were distributed into 22 cell clusters (SI Appendix, Fig. S1A). Expressions of human cytokeratin 7 (*KRT7*) and human leukocyte antigen-A (*HLA-A*) distinguished the trophoblast cell clusters from non-trophoblast cells (SI Appendix, Fig. S1B). As GATA3 and TFAP2C are expressed in all mononuclear trophoblast cells of a first-trimester human placenta (31), we tested and confirmed expressions of *GATA3* and *TFAP2C* mRNA in all single cell clusters comprising trophoblast cells (SI Appendix, Fig. S1C).

To identify distinct CTB subpopulations in single cell clusters, we compared expressions of *ELF5, CCNB2, ITGA2, ERVFRD1* and *HLA-G* (Fig. 1C). *ELF5,* which is expressed in the CTB subpopulation that maintains an undifferentiated stem-state, was detected mainly in cells of clusters 1 and 12 (Fig. 1C, upper left panel) and within a few cells of cluster 4. A recent study showed that the stem-state CTBs also express Basal Cell Adhesion Molecule (BCAM) (17) and analyses of single cell clusters showed that *BCAM* is predominantly expressed in *ELF5* expressing cells of clusters 1, 12 and cluster 4 (SI appendix, Fig. S1D). Thus, we concluded that the *ELF5*/*BCAM*-expressing cells within clusters 1,12 and 4 comprise the undifferentiated stem-state CTB subpopulation. Expression of *CCNB2*, which marks all mitotically active CTBs of a first-trimester placenta, was detected in all *ELF5*-expressing cells of clusters 1, 12 and 4. *CCNB2* expression was also detected in a majority of cells within clusters 3-6 and 14 and in a few cells of cluster 2 (Fig. 1C, upper middle panel), indicating that those cell clusters contain mitotically active but differentiating CTBs. Interestingly, mRNA expression of *ITGA2*, which marks the CTB progenitors at the proximal column of an anchoring villous, was not detected in *ELF5*-expressing CTBs within clusters 1 and 12. Rather, *ITGA2* expressing cells were scattered within cells of cluster 2, 3, 4, 5, 6, and 14 (Fig. 1C, upper right panel). We also noticed a similar expression pattern of *NOTCH1* (SI appendix, Fig. S1D). Thus, we concluded that the *ITGA2*-expressing cells of cluster 2, 3, 4, 5, 6 and 14 represent the proximal column CTB progenitors within anchoring villi.

*ERVFRD1,* an endogenous retroviral encoded cell-fusion gene, is shown to be expressed in mitotically inactive CTBs within a first-trimester human placenta (16). We detected *ERVFRD1* mRNA expression only within single-cells of cluster 11 (Fig. 1C, lower left panel), indicating that cells in cluster 11 represent the mitotically inactive CTBs that are committed for STB differentiation. High-level expression of HLA-G, which is induced in developing EVTs, was detected within cells of clusters 9, 18, 10 and a few cells of cluster 2 (Fig. 1C, lower middle panel). Interestingly, the high level *HLA-G* expressing cells of clusters 9, 18, 10 and 2 did not express ITGA2. Thus, we concluded that those cells were committed for EVT differentiation.

Next, we tested expressions of Hippo signaling cofactors in single cell clusters, representing different CTB subpopulations of a first-trimester human placenta. We found that *VGLL1* is highly expressed in almost all single trophoblast cells within a first-trimester placenta (Fig. 1C). However, *WWTR1* and *YAP1* showed contrasting expression patterns in *ELF5*-expressing stem-state CTBs vs. *CCNB2*-expressing differentiating CTB progenitors (Fig. 1D). *WWTR1* and *YAP1* are expressed in both stem-state and differentiating CTBs of clusters 1, 12 and 4, although, *YAP1* expression is higher in stem-state CTBs. In contrast, *WWTR1* expression is induced in differentiating CTBs of cell clusters 2, 3, 4, 5, 6 and 14 (Fig. 1D). Cell clusters 2, 3, 5, 6 and 14 also contained *ITGA2-*expressing column CTB progenitors. We validated *WWTR1* expression in nearly all *ITGA2*-expressing CTBs, whereas *YAP1* expression was detected in a small fraction of *ITGA2*-expressing cells (SI Appendix, Fig. S2A, B). Interestingly, a contrasting expression pattern of *YAP1* and *WWTR1* was also observed in mitotically inactive CTBs vs. emerging EVTs. YAP1 expression is maintained in *ERVFRD1-*expressing, mitotically inactive CTBs of cluster 11, and is reduced in *HLA-G* expressing cells of clusters 9, 18, 10 and 2 (Fig. 1D). In contrast, *WWTR1* is highly expressed in *HLA-G* expressing emerging EVTs and is repressed in mitotically arrested CTBs (Fig. 1D).

Next, we tested WWTR1 expression in human TSCs that were derived from first-trimester CTBs. Reverse transcription followed by quantitative PCR (RT-qPCR) showed that *WWTR1* mRNA is expressed in undifferentiated human TSCs and the expression is induced during EVT differentiation (SI Appendix, Fig. S3A). Western blot analyses confirmed induction of WWTR1 protein expression during EVT differentiation in human TSCs (Fig. 1E). Immunofluorescence analyses showed that WWTR1 is localized within nuclei in both undifferentiated TSCs and in TSC-derived EVTs (SI Appendix, Fig. S3B). Collectively, our expression analyses indicated that during human placentation WWTR1 might have important functional roles in CTB progenitors and promote their differentiation to EVTs.

### WWTR1 regulates self-renewal of human trophoblast progenitors

WWTR1 is expressed in mitotically active CTBs within a first-trimester placenta and expression is maintained in CTB-derived Human TSCs. Therefore, we performed loss-of-function analyses to test the importance of WWTR1 in human trophoblast progenitor self-renewal. We depleted WWTR1 in human TSCs (*WWTR1-KD* human TSC) by RNA interference (RNAi) using lentiviral-mediated transduction of small hairpin RNAs (shRNAs) (Fig. 2A, B). In a culture condition that promotes human TSC proliferation at stem-state, *WWTR1-KD* human TSC showed loss of stem-state colony morphology and strong reduction in cell proliferation ability (Fig. 2C-E). We also tested self-renewal of *WWTR1-KD* human TSC by assessing their ability to form self-renewing 3-dimensional trophoblast organoids (TSC organoids). Unlike the control human TSCs, *WWTR1*-KD human TSCs showed severe impairment in organoid formation (Fig. 2F). Control human TSCs formed large organoids with prolonged culture (8-10 days) and could be dissociated and reorganized to form secondary organoids, indicating the self-renewing ability. In contrast, *WWTR1*-KD human TSCs formed much smaller organoids, which were not maintained upon passaging.

**Figure 2:**
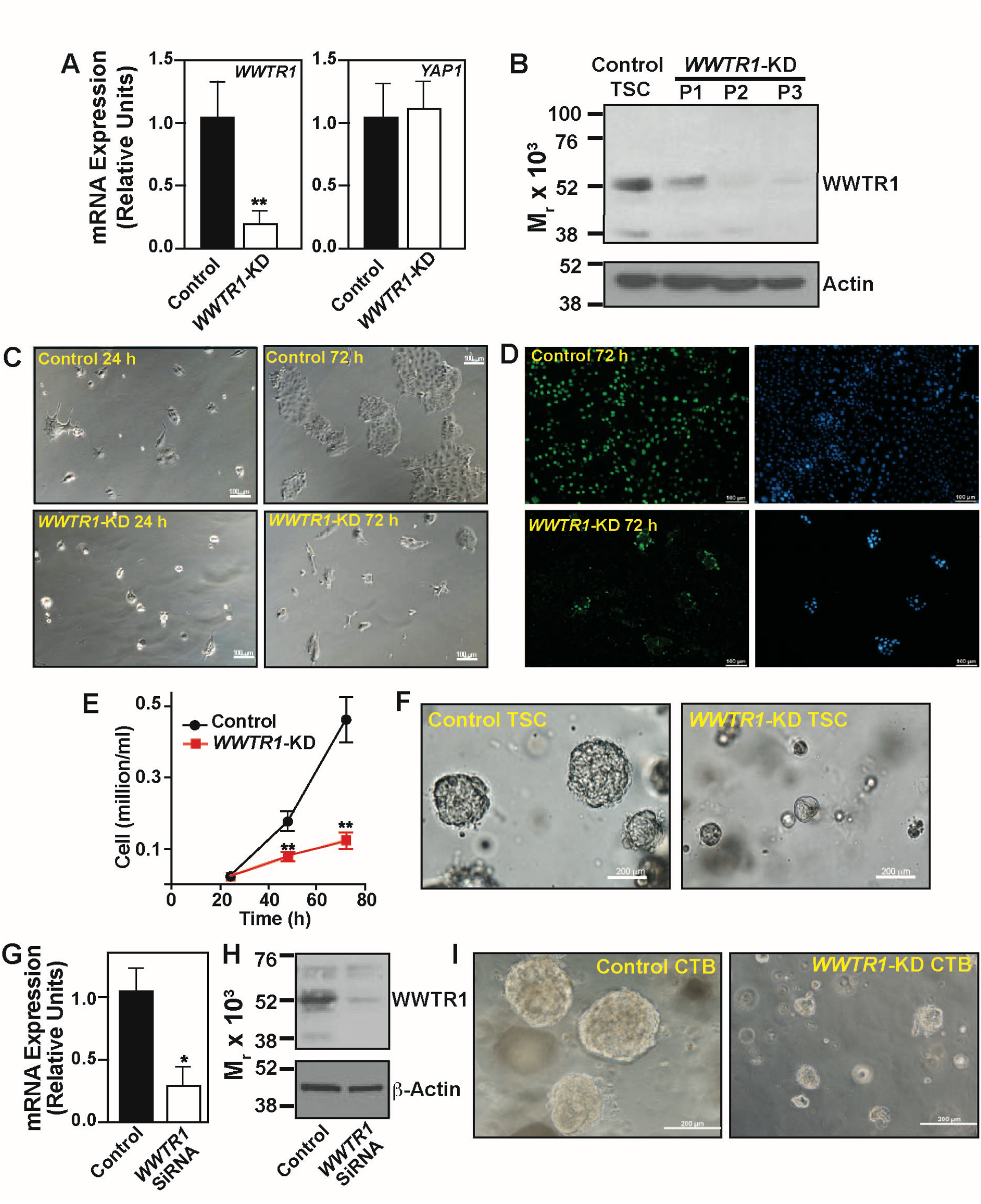
Loss-of WWTR1 impairs self-renewal in human TSCs and primary CTBs. (A) *WWTR1* and *YAP1* mRNA expressions were tested in human TSCs without (control) and with WWTR1-depletion by shRNA [*WWTR1-*KD]. Plot shows strong reduction of *WWTR1-*mRNA expression in *WWTR1-*KD human TSCs did not significantly alter *YAP1* mRNA expression (mean ± SE; n = 3, p≤0.005) (B) Western blot analyses showing depletion of WWTR1 protein expression in *WWTR1-*KD human TSCs over three passages. (C) Equal number of control and *WWTR1*-KD human TSCs were plated and cultured in stem-state culture condition. Micrographs confirm reduced cell proliferation in *WWTR1*-KD human TSCs. (D) Images show BRDU incorportaion in control and WWTR1-depleted human TSCs, when cultured in stem-state culture condition over 72h. (E) Plot shows growth kinetics of human TSCs, without and with WWTR1 depletion. (F) Micrographs show inefficient organoid formation by *WWTR1*-KD human TSCs. (G) and (H) RT-qPCR and Western blot analysis, respectively, showing RNAi-mediated depletion of WWTR1 expression in primary CTBs, isolated from human first-trimester placentae. (I) Micrographs show inefficient organoid formation by WWTR1-depleted (*WWTR1*-KD) primary CTBs.

Next, we tested the impact of WWTR1 depletion on the self-renewal ability of primary CTBs that were isolated from first-trimester (6-10 weeks) human placentae. We depleted WWTR1 expression in primary CTBs via small interfering RNA (SiRNA) molecules (Fig. 2G, H) and tested the ability to form self-renewing 3-dimensional CTB-organoids. We found that similar to human TSCs, WWTR1-depletion in primary CTBs strongly inhibited organoid formation efficiency (Fig. 2I). Thus, loss-of-function studies in human TSCs and primary CTBs strongly indicated that WWTR1 plays an important role to maintain the self-renewal ability within mitotically active CTB progenitors of developing human placenta.

### WWTR1 directly regulates TP63 expression in human trophoblast progenitors

To understand how WWTR1 regulates trophoblast progenitor self-renewal, we performed global gene expression (RNA-seq) analyses in *WWTR1*-KD human TSCs. Depletion of WWTR1 in human TSCs significantly altered expression of 960 genes (216 down-regulated and 744 up-regulated, SI Appendix, Dataset S1, Fig. S4A). RNA-seq analyses revealed that mRNA expression of *TP63*, which is implicated in maintenance of CTB stem-state (32), is strongly down-regulated in *WWTR1*-KD TSCs (Fig. 3A, SI Appendix, Fig. S4B). We confirmed TP63 down-regulation in *WWTR1*-KD TSCs via RT-qPCR and immunofluorescence analyses (Fig. 3B, C). Furthermore, using quantitative chromatin immunoprecipitation (ChIP-PCR) we detected WWTR1 occupancy at a conserved TEAD motif at the *TP63* locus in undifferentiated TSCs (Fig. 3D, E). These results indicate that WWTR1-mediated induction of *TP63* expression might be one of the molecular mechanisms to maintain self-renewal in stem-state CTBs within a developing human placenta.

**Figure 3.**
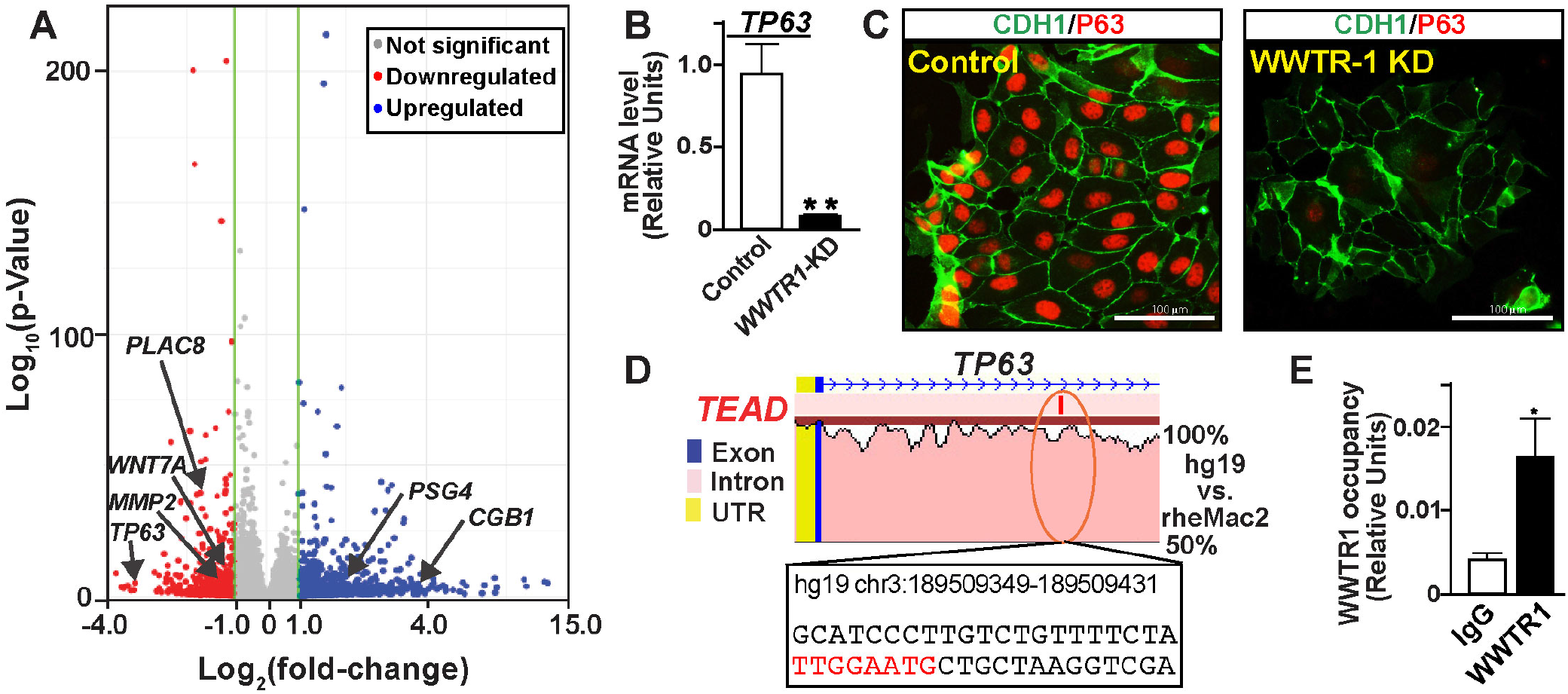
WWTR1 directly regulates TP63 expression in human TSCs. (A) Volcano plot showing global gene expression changes in *WWTR1*-KD human TSCs. Unbiased RNA-seq analyses were performed and ≥2-fold gene-expression changes in *WWTR1*-KD human TSCs with a false discovery rate of P < 0.05 were indicated with colored dots (blue: up-regulated, red: down-regulated). Significant down-regulation in expression of *TP63 (*a marker of undifferentiated CTBs*), PLAC8, MMP2 (*important for EVT development*), WNT7A,* and up-regulation of *CGB1* and *PSG*4 (STB-specific genes) are indicated. (B) RT-qPCR (mean ± SE; n = 3, p≤0.001) and (C) Immunofluorescence images, respectively, confirming down-regulation of TP63 expression in *WWTR1-*KD human TSCs. (D) rVISTA alignment plot of a conserved TEAD-motif containing region of human and rhesus macaque *TP63* genes. The conserved TEAD-motif (in red letters) along with adjacent base sequences and associated coordinate on the human *TP63* gene are indicated. (E) The plot shows WWTR1 occupancy at the conserved TEAD-motif of the *TP63* locus (shown in panel D) in human TSCs (mean ± SE; n = 3, p≤0.01).

### Discovery of a WWTR1-WNT regulatory axis in human trophoblast progenitors

Unbiased Gene Set Enrichment Analysis (GSEA) of RNA-seq data showed that loss of WWTR1 in human TSCs down-regulates transcription of various genes in the Wingless/Integrate (WNT) signaling pathway (Fig. 4A). A detailed look at the expression of WNT genes showed that six WNT genes, *WNT3, WNT4, WNT5B, WNT7A, WNT8B* and *WNT9A*, are repressed in *WWTR1*-KD TSCs (Fig. 4B). The WNT signaling pathway has been implicated as a key regulator for maintaining CTBs at a progenitor state (26). Gene expression analyses in first-trimester CTBs showed that many of the WNT genes, including WNT3 and WNT7A, are expressed in undifferentiated CTBs (25) and activation of WNT signaling was key to successful derivation of human TSCs (25) and self-renewing CTB organoids (26, 27). Therefore, we looked at expressions of WNT genes in CTB progenitors of a first-trimester human placenta at single-cell resolution.

**Figure 4.**
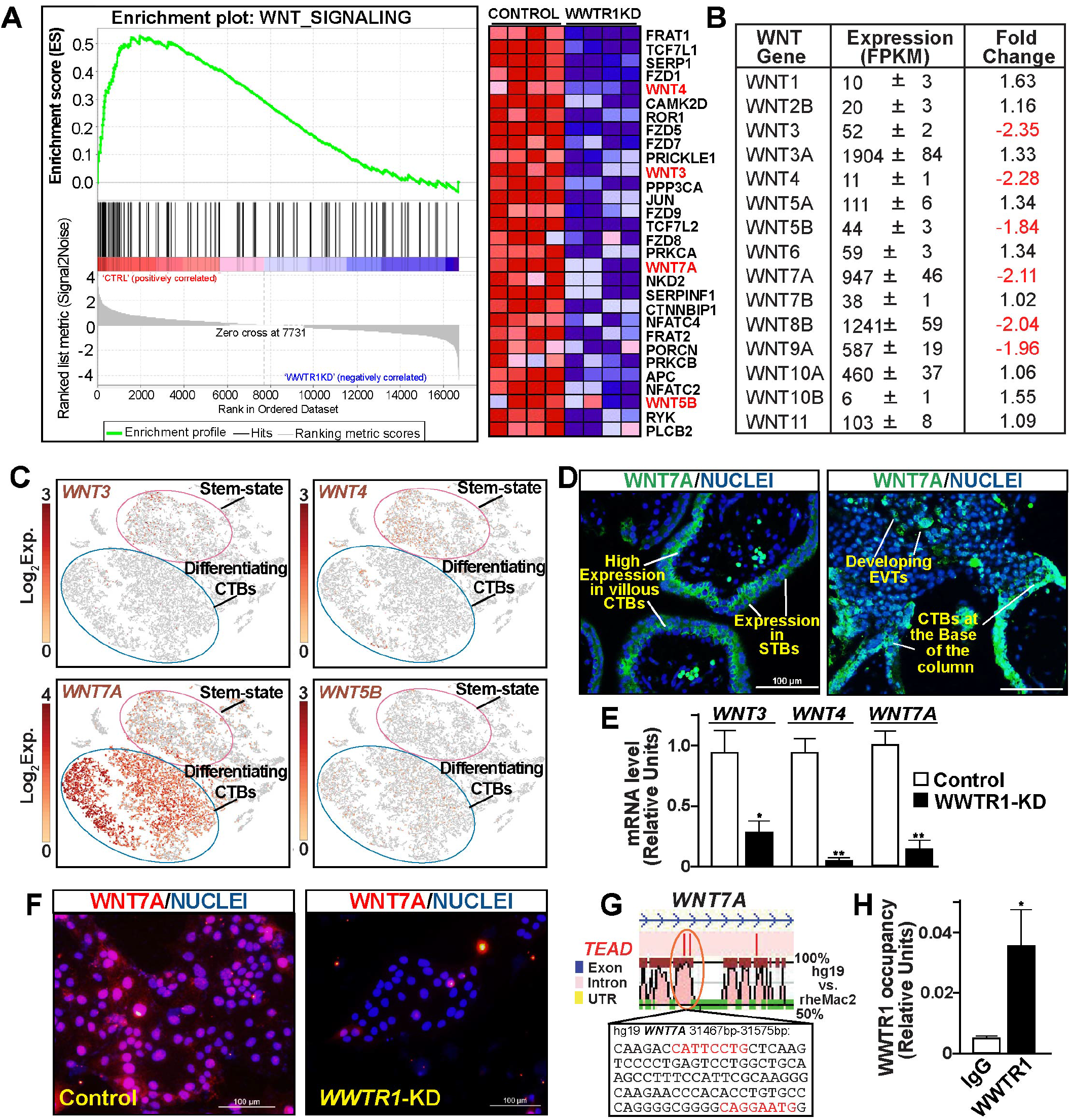
WWTR1 regulates WNT signaling components in human TSCs. (A) GSEA of differentially expressed genes showing down-regulation of WNT-signaling genes in *WWTR1-*KD human TSCs. The heat map shows specific genes that were down-regulated in *WWTR1*-KD human TSCs. Various WNT family members are highlighted. (B) The table shows average expression levels of all *WNT* genes that were expressed in human TSCs. WNT Genes that were down-regulated in *WWTR1*-KD human TSCs are highlighted. (C) t-SNE plots showing differential mRNA expression patterns of *WNT* genes in single cell clusters representing stem-state and differentiating CTBs of first-trimester human placentae. (D) Immunofluorescence images show *WNT7A* protein expression in trophoblast cells of first-trimester placental villi. Representative images of floating and anchoring villi are shown. (E) RT-qPCR analyses confirming *WNT3, WNT4* and *WNT7A* mRNA expressions in *WWTR1-*KD human TSCs (mean ± SE; n = 4, *; p<0.01, **; p≤0.001). (F) Immunofluorescences images confirms loss of and WNT7A protein expression in *WWTR1-*KD human TSCs. (G) rVISTA alignment plot of a conserved TEAD-motifs containing region of human and rhesus macaque *WNT7A* genes. Conserved TEAD-motif (in red letters) along with adjacent base sequences and associated coordinate on the human *WNT7A* gene, where WWTR1 occupancy was detected in human TSCs, are indicated. (H) Quantitative ChIP analysis identified WWTR1 occupancy at the region with highlighted conserved TEAD-motifs of the *WNT7A* locus (shown in panel G) in human TSCs (mean ± SE; n = 3, p<0.01).

scRNA-seq analyses showed that among six WNT genes, namely *WNT3, WNT4, WNT5B, WNT7A, WNT8B* and WNT9A, which are regulated by WWTR1 in human TSCs, *WNT7A* is most abundantly and widely expressed in mitotically active CTBs (Fig. 4C). *WNT3* and *WNT4* are also expressed in undifferentiated stem-state CTBs. However, compared to *WNT7A*, *WNT3* and *WNT4-*expressing CTBs are less abundant in a first-trimester placenta (Fig. 4C). *WNT5B* mRNA is not expressed in undifferentiated CTBs; rather, *WNT5B* mRNA expression was detected in a small number of differentiating, mitotically active CTBs (Fig. 4C), and *WNT8B* and *WNT9A* are not expressed in primary CTBs (SI Appendix, Fig. S5).

As scRNA-seq analyses identified WNT7A as the most abundantly expressed WNT gene in first-trimester CTBs, we tested WNT7A protein expression in human first-trimester placenta. We found that WNT7A is highly expressed in CTBs within floating villi, and expression is reduced but maintained in STBs (Fig. 4D, left panel). In anchoring villi, WNT7A is also expressed at the base of the CTB column and in the emerging EVTs at the distal cell column (Fig. 4D, Right Panel).

Our expression analyses showed that among WWTR1-regulated WNT genes in human TSCs, WNT3, WNT4 and WNT7A are expressed in primary CTBs and may contribute to the CTB self-renewal process. Therefore, we validated down-regulation of *WNT3, WNT4* and *WNT7A* mRNA expression in *WWTR1*-KD TSCs via RT-qPCR (Fig. 4E). We also validated loss of WNT7A protein expression in WWTR1-KD TSCs (Fig. 4F). Furthermore, using ChIP-PCR we found that WNT7A is a direct WWTR1 target gene in human TSCs (Fig. 4G, H). Collectively, our experiments identified a WWTR1-WNT regulatory axis that could orchestrate gene expression programs in human CTB progenitors.

### WWTR1 prevents induction of STB fate and promotes EVT differentiation in human trophoblast progenitors

Global gene expression analyses in *WWTR1*-KD human TSCs revealed strong up-regulation of many STB-specific genes, such as chorionic gonadotropin A (CGA), chorionic gonadotropin B isoforms (CGBs) and pregnancy-specific beta-1-glycoproteins (PSGs), in a culture condition that maintains human TSC stem-state (Fig. 3A, SI Appendix, Dataset S1 and Fig. S4A, B). We confirmed induction of STB-specific gene transcripts in *WWTR1*-KD human TSCs via RT-qPCR (Fig. 5A). We also found that siRNA-mediated depletion of WWTR1 in primary CTBs of first-trimester human placentae strongly induced CGB protein expression and secretion (Fig. 5B, C). Furthermore, at human TSC stem-state culture condition, extended culture of *WWTR1*-KD human TSCs often resulted in spontaneous cell-fusion and formation of multinucleated syncytium (Fig. 5D). The nuclei of those multinucleated syncytium expressed high levels of CGB, confirming induction of STB-differentiation fate in *WWTR1*-KD human TSCs in a culture condition that should maintain the TSC stem state. Taken together, our studies strongly indicated that, during human placentation, WWTR1 function in CTB progenitors prevents induction of the STB differentiation program by suppressing expression of STB-specific genes.

**Figure 5.**
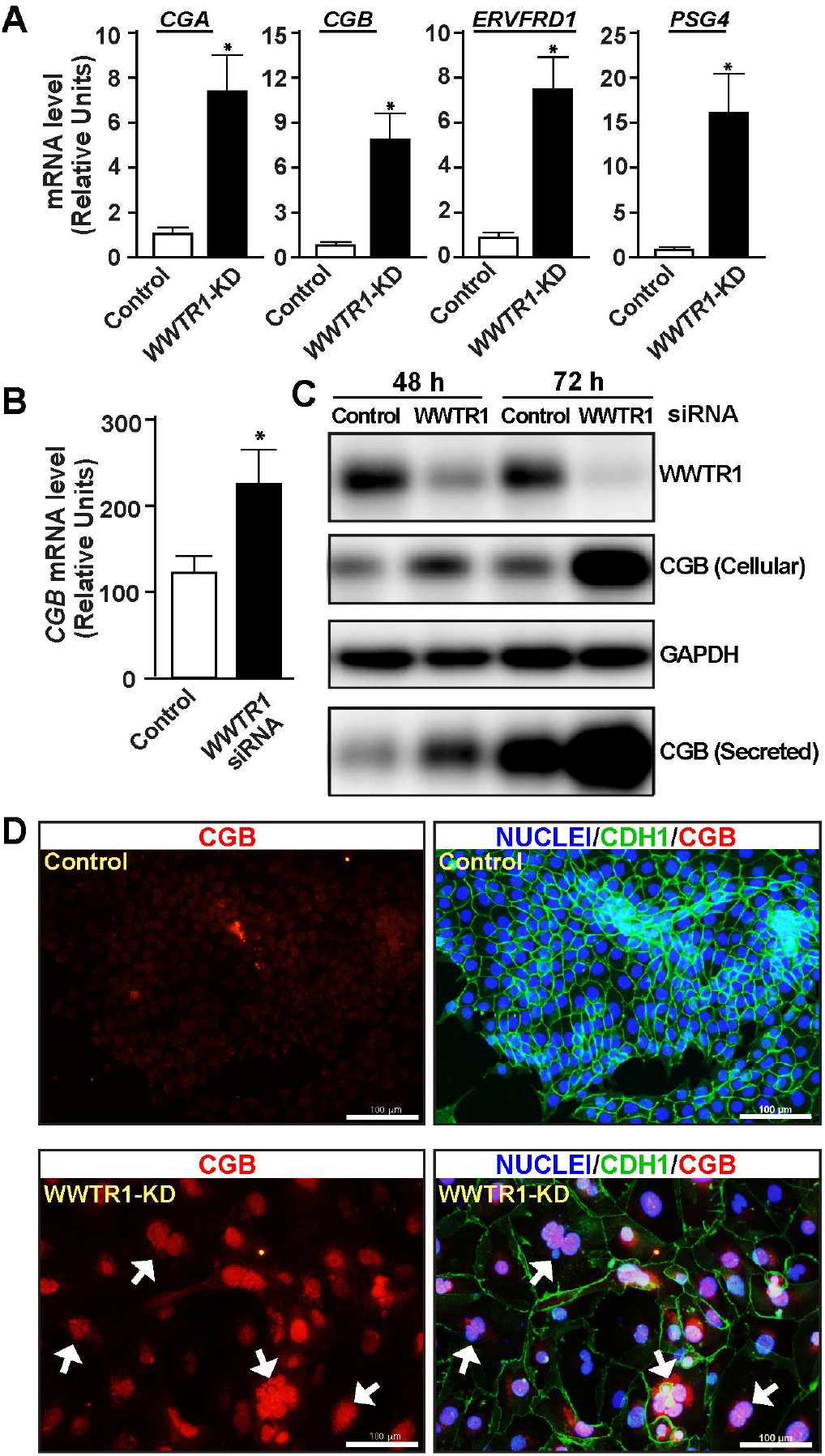
WWTR1 prevents STB-differentiation in human TSCs and primary CTBs. (A) RT-qPCR analyses confirming induction of mRNA expressions STB-specific genes*, CGA, CGB, ERVFRD1* and *PSG4* in *WWTR1-*KD human TSCs (mean ± SE; n = 4, p≤0.005). (B) RT-qPCR analyses confirming *CGB* mRNA expressions in CTBs, isolated from first-trimester human placentae upon siRNA-mediated depletion of *WWTR1* (mean ± SE; n = 3, p≤0.01). (C) Western blot analyses confirming induction of CGB protein expression in *WWTR1-*depleted CTBs (D) Fluorescence images show propensity of STB-differentiation in *WWTR1-*KD human TSCs, when cultured in stem-state culture conditions for three passages. *WWTR1-*KD human TSCs showed enhanced propensity of cell fusion with multi-nucleated cells and loss of E-Cadherin expression (white arrows). The nuclei of fused cells also showed higher level of CGB protein expression.

The global gene expression analyses also revealed that mRNA expressions of *matrix metalloproteinase 2* (*MMP2*), *Placenta Associated 8* (*PLAC8*) and *SMAD Family Member 3* (*SMAD3*), which are implicated in EVT development (33-35), were down-regulated in *WWTR1*-KD human TSCs (SI Appendix, Dataset S1). MMP2 and other MMPs have been implicated in EVT development and invasion (33, 36). Thus, we tested mRNA expressions of MMP family members along with *PLAC8* and *SMAD3* in *WWTR1*-KD TSCs using RT-qPCR. We found that along with *PLAC8* and *SMAD3,* mRNA expressions of four MMP genes, *MMP2, MMP11, MMP14* and *MMP15* were significantly down-regulated in *WWTR1*-KD human TSCs (Fig. 6A). We also confirmed loss of MMP2 and SMAD3 protein expressions in *WWTR1*-KD human TSCs (Fig. 6B). Our single-cell gene expression analyses in first-trimester human placenta showed that all of these genes are either highly expressed (*MMP2*, *MMP14*, and *PLAC8*) or highly induced (*MMP11*, *MMP15* and *SMAD3*) in developing EVTs (cell clusters 9, 18, 10 and 2; Fig 6C, SI Appendix Fig. S6A), in which HLA-G expression was also induced (shown in SI Appendix, Fig. S1C). Therefore, we next tested the importance of WWTR1 in EVT development.

**Figure 6.**
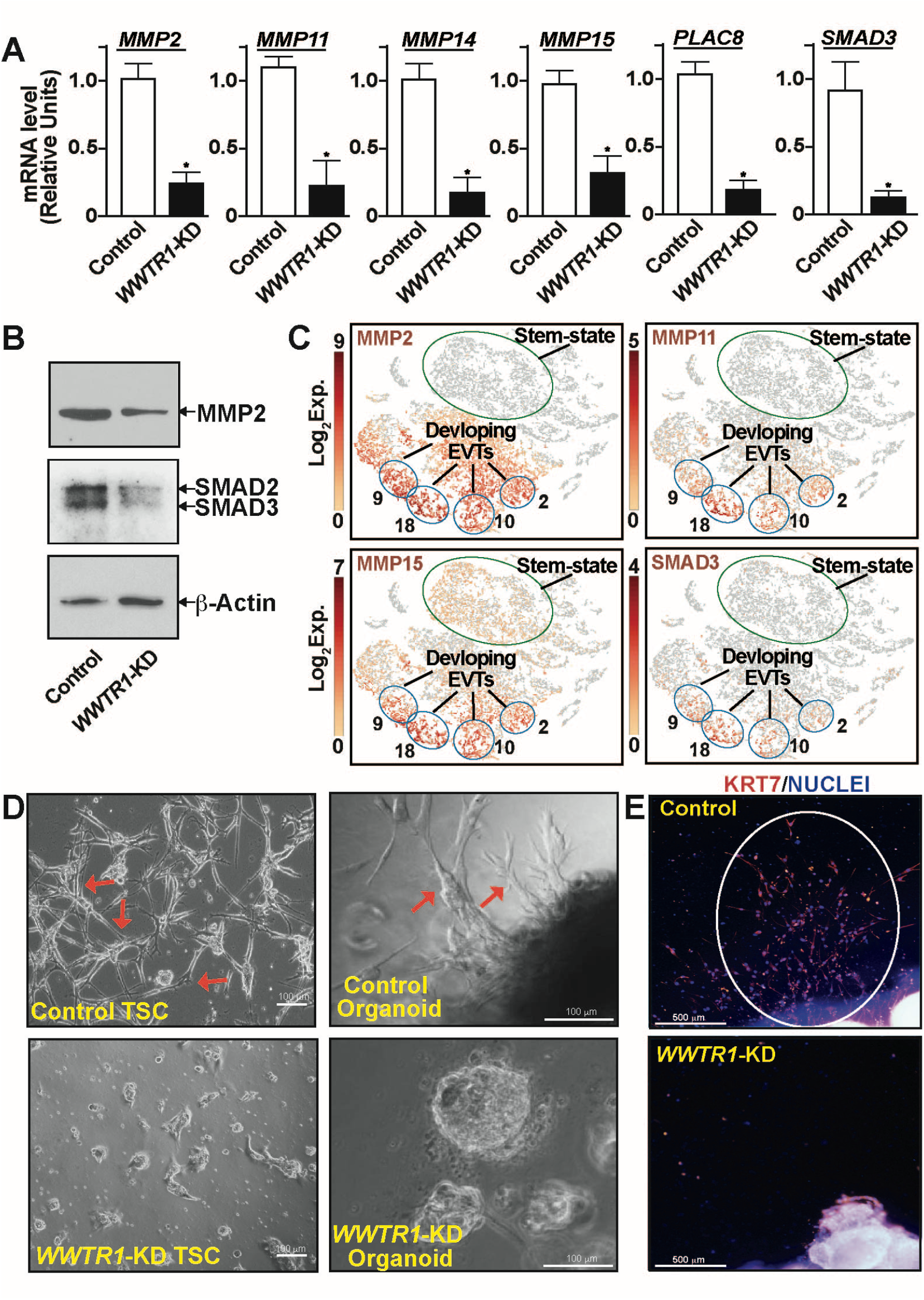
WWTR1 regulates EVT development. (A) RT-qPCR analyses confirming down-regulation of mRNA expressions of *MMP2, MMP11, MMP14, MMP15, PLAC8* and *SMAD3* in *WWTR1-*KD human TSCs (mean ± SE; n = 4, p<0.01). (B) Western blot analyses confirming loss of MMP2 and SMAd3 protein expressions in *WWTR1-KD* human TSCs. (C) t-SNE plots showing mRNA induction of *MMP2, MMP11, MMP15* and *SMAD3* in single cell clusters representing developing EVTs of first-trimester human placentae. Note that the cell clusters representing stem-state CTBs mostly lack mRNA expressions of *MMP2, MMP11,* and *SMAD3* and have much less mRNA expression of *MMP15*. (D) Representative phase contrast images show inefficient EVT development from *WWTR1-*KD human TSCs (left panels) and first-trimester CTB-Organoids (right panels). In a culture condition that promote EVT differentiation, control TSCs and CTB-organoids readily developed EVTs with characteristic elongated spindle-shaped cell protrusions (shown in red arrows). However, EVT development was strongly impaired from *WWTR1-*KD human TSCs and *WWTR1-*KD CTB organoids. (E) Immunofluorescence images show impairment of EVT emergence from human first-trimester placental explants upon WWTR1-depletion. Invasive EVTs were readily developed (highlighted with white ellipse) when first-trimester placental explants were cultured on matrigel in a culture condition that promote EVT differentiation. EVT emergence was strongly inhibited from placental explants, in which *WWTR1* expression was depleted.

We performed three different experiments to test the importance of WWTR1 in EVT development. First, we tested EVT differentiation efficiency of *WWTR1*-KD human TSCs and found that loss of WWTR1 in human TSC strongly inhibits the efficiency of EVT differentiation (Fig. 6D, left panels). Next, we studied first-trimester CTB-derived organoids, which has been successfully utilized to test EVT development from primary CTBs (26, 27). EVT development was readily noticed when control CTB-organoids were cultured on matrigel. However, RNAi-mediated silencing of WWTR1 expression nearly abrogated EVT emergence from CTB-organoid (Fig. 6D, right panels). Finally, we tested EVT emergence from human first-trimester placental explants after depleting WWTR1 expression via RNAi (SI Appendix, Fig. S6B-C). Similar to our findings with human TSCs and primary CTBs, EVT emergence from first-trimester placental explants was strongly inhibited upon depletion of WWTR1 expression (Fig. 6E and SI Appendix, Fig. S6D). Collectively, our studies in human TSCs, primary CTBs and placental explants identified WWTR1 as an important regulator of EVT development.

### Extreme Preterm birth is associated with loss of WWTR1 expression in CTBs

Defective trophoblast development has been implicated as a major cause of pregnancy-associated diseases, including preterm birth, IUGR and preeclampsia. It was shown that extreme preterm birth is often associated with premature differentiation of villous CTBs (37). Furthermore, pregnancies associated with severe PE or severe IUGR often demonstrate depletion of proliferating CTBs (38). In addition, severe PE is also associated with increased syncytial knot formation (39), indicating that these pregnancies are associated with an imbalance in CTB self-renewal vs. differentiation process. As we discovered WWTR1 as one of the important regulators to maintain self-renewal ability in CTBs, we tested whether *WWTR1* mRNA expression was altered in placentae from pregnancies that are associated with preterm birth, IUGR and PE (Fig. 7A, B).

**Figure 7.**
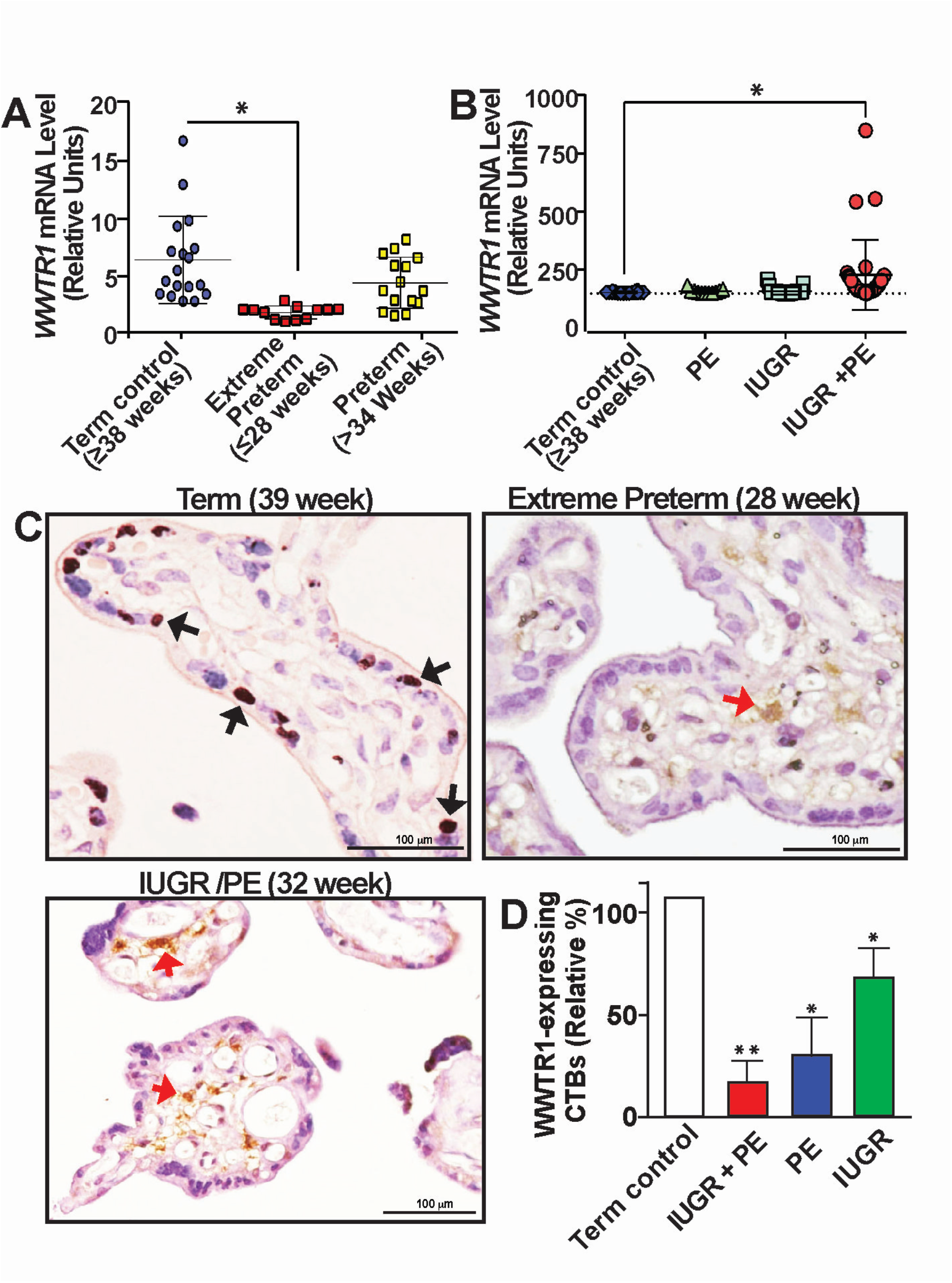
WWTR1 expression is impaired in CTBs in pathological pregnancies. (A) RT-qPCR analyses of *WWTR1* mRNA expression from mRNAs isolated from whole placental tissues from pregnancies with gestational age ≥38 week (Term control), preterm-birth (≥34 weeks) or extreme preterm birth (≤28 weeks). (* indicates significant change (p<0.01) in *WWTR1* mRNA expression in placentae from extreme preterm birth) (B) RT-qPCR analyses of *WWTR1* mRNA expression from mRNAs isolated from whole placentae that were collected from pregnancies with preterm birth along with IUGR, PE or both IUGR and PE (IUGR/PE). (* Indicates significant change (p<0.01) in *WWTR1* mRNA expression in placentae with preterm birth along with IUGR/PE)). (C) Representative immunostained images show lack of WWTR1-expressing CTBs (Black arrows) in placentae from pregnancies that are associated with extreme preterm birth or preterm birth in association with IUGR/PE. Red arrows indicate WWTR1-expressing non-trophoblast cells. (D) WWTR1 expressing CTBs were quantitated from 10 different placental sections from normal term pregnancy or from preterm pregnancies with IUGR, PE or IUGR/PE. The plot shows significant (p<0.01) reduction in WWTR-1 expressing CTBs in pathological pregnancies.

First, we tested WWTR1 expression in placentae, which were associated with preterm birth without any reported complications of IUGR and PE, and compared that with placentae that are associated with normal term pregnancy. We analyzed 27 placentae from preterm pregnancies without IUGR and PE, of which 12 placentae were associated with extreme preterm birth (babies were born at or before 28 weeks of pregnancy). Intriguingly, we found that WWTR1 mRNA expression levels were significantly reduced in placentae from pregnancies associated with extreme preterm birth (Fig. 7A). However, we have not noticed any significant change in WWTR1 mRNA expression within placentae from preterm births, in which babies were born after 34 weeks of pregnancy. As WWTR1 is predominantly expressed in CTBs within a term placenta, we tested WWTR1 protein expressions in placental sections associated with extreme preterm birth and found that the number of WWTR1 expressing CTBs are drastically reduced in placentae from extreme preterm birth (Fig. 7C).

As IUGR and PE are often associated with preterm birth, we also tested *WWTR1* mRNA expression in placentae from preterm pregnancies, in which pregnancy duration was less than 34 weeks and were also characterized with either IUGR or PE or IUGR in combination with PE (IUGR/PE). We analyzed 44 IUGR placentae, 11 PE placentae and 31 IUGR/PE placentae with average pregnancy duration of 33 weeks, 32 weeks and 30 weeks, respectively. We found that *WWTR1* mRNA expressions in whole placentae are not significantly altered in pregnancies associated with IUGR or PE, whereas placentae associated with IUGR/PE pregnancies showed induction of WWTR1 mRNA expression in whole placenta (Fig. 7B). However, when we tested WWTR1 protein expressions via immunostaining, we hardly noticed presence of WWTR1-expressing CTBs in placentae from pregnancies with preterm birth along with IUGR/PE (Fig. 7C, 7D and SI Appendix, Fig. S7). Significant reduction in WWTR1-expressing CTBs was also noticed in placentae from preterm pregnancies with IUGR and PE (Fig. 7D and SI Appendix, Fig. S7). In contrast, we noticed increased infiltration of WWTR1-expressing cells within the stroma of those placentae (Red arrows, Fig. 7C). These results indicated that the increased WWTR1 mRNA expression that we observed with whole placental tissues from IUGR/PE were not due to increased expression in CTBs or STBs. Rather; preterm birth with IUGR/PE is often associated with loss of WWTR1-expressing CTB population in placental villi. Taken together, our findings indicated that impaired WWTR1 function in CTB progenitors could be one of the trophoblast-associated molecular causes in pathological pregnancies, such as extreme preterm birth.

## DISCUSSION

An important aspect of human placentation is establishment of distinct gene expression programs in CTB progenitors of anchoring vs. floating villi. During early stages of human placentation, gene expression programs in floating villi are fine-tuned to support self-renewal of CTB progenitors as well as instigation of the STB-specific differentiation program. The proper balance of CTB self-renewal and STB differentiation in floating villi ensure development of a mature placenta with an enormous surface area for nutrient and gas exchange at the maternal-fetal interface. Furthermore, continuous shedding of apoptotic STBs and incorporation of new STBs from the underlying CTB layer ensure maintenance of the functional STB layer throughout pregnancy. In contrast, in anchoring villi, gene expression programs within CTB progenitors are orchestrated to promote extensive proliferation to form a CTB column and to instigate an EVT-specific differentiation program in the distal cells of the CTB column. This extensive proliferation of the CTB column and EVT differentiation predominantly takes place during the first trimester of pregnancy. Thus, success in human pregnancy relies on the establishment of proper spatial and temporal gene expression programs in CTB progenitors of a developing placenta. Our findings in this study establish the hippo signaling cofactor WWTR1 as an essential regulator to orchestrate the gene expression program and balance self-renewal vs. differentiation in CTB progenitors. Our experimental findings indicate a bimodal function of WWTR1 in human trophoblast progenitors. In floating villi, it promotes CTB self-renewal and suppresses premature instigation of the STB differentiation fate, whereas in anchoring villi, WWTR1 function in CTB progenitors is important to instigate EVT differentiation. We also discovered that pregnancies associated with extreme preterm birth as well as IUGR/PE are often characterized with loss of WWTR1 expression in CTBs. Collectively, our findings implicate defective WWTR1 expression/function in CTBs as one of the molecular causes for adverse pregnancies.

Our findings that WWTR1 promotes CTB self-renewal along with our earlier reports showing essential roles of TEAD4 and YAP1 in maintenance of CTB self-renewal establishes the critical importance of the Hippo signaling pathway in human trophoblast development. These findings also indicate a functional redundancy of WWTR1 and YAP1 in CTB progenitors. However, our single cell resolution gene expression analyses indicated that YAP1 and WWTR1 are differentially expressed in distinct CTB subpopulations. High level of YAP1 expression is confined within the ELF5-expressing undifferentiated/stem-state CTB progenitors. Interestingly, expression of TEAD4 was also predominantly detected within the ELF5-expressing CTB subpopulations (11). In contrast, WWTR1 expression is induced in the CTB sub-population, which are mitotically active but poised for differentiation. Thus, we propose that a TEAD4/YAP1 transcriptional complex is important to maintain a ground level of stemness within undifferentiated CTB progenitors, whereas WWTR1 can interact with other TEAD family members to maintain the self-renewal ability in CTB progenitors, which are priming for differentiation, including the column CTBs of anchoring villi. Future studies involving identification of global targets in CTBs along with spatial single-cell genomics of developing human placenta will be instrumental to gain insights into the transcriptional programs that are established by YAP1 and WWTR1 in distinct CTB sub-populations.

Crosstalk of WNT signaling components with WWTR1 and YAP1 has been identified as an important regulatory axis in several cellular systems (40). The WNT signaling pathway has also been implicated in the CTB self-renewal process (26) as well as their differentiation to EVTs (41). However, the roles of individual WNT molecules in maintaining CTB self-renewal vs. differentiation are not well understood. Our findings in this study indicate a prominent role of WNT7A in human trophoblast development. We discovered WNT7A as the most abundantly expressed WNT molecule in CTBs of a first-trimester placenta, and its expression pattern is similar to WWTR1 in subpopulations of mitotically active CTBs and is a direct target of WWTR1 in CTB-derived human TSCs. WNT7A is highly expressed in CTBs within floating villi and at the base of the CTB column in anchoring villi. WNT7A expression is also detected in emerging EVTs at the distal columns of anchoring villi, indicating that the WWTR1-WNT7A signaling axis could be important for the CTB self-renewal process within the floating villi and EVT development in anchoring villi. WWTR1 also regulates expressions of other WNT molecules, such as WNT3 and WNT4 in human TSCs. Interestingly, our scRNA-seq analyses showed that, in a first-trimester human placenta, WNT3 and WNT4 expressions, albeit at low level, are confined in stem-state CTBs, which also express high-levels of YAP1 and TEAD4. Thus, crosstalk among distinct WNT molecules and Hippo-signaling components might regulate gene expressions in different CTB sub-populations within a developing human placenta.

We found that WWTR1 is essential for EVT differentiation in human TSCs and emergence of EVT cells from first-trimester placental explants. Interestingly, WWTR1 function in EVT development appears to be distinct from YAP1, which is suppressed in EVTs. We have shown that during EVT differentiation of human TSCs, WWTR1 is required for optimal expression of MMP2, MMP11, MMP14 and MMP15. MMP2 has been shown to regulate EVT invasion (33, 36). Expressions of MMP11, MMP14 and MMP15 were earlier detected in human trophoblast cells, including EVTs within maternal decidua (36, 42). An earlier study showed that PLAC8 is selectively induced during EVT development and induces the formation of filopodia in migratory trophoblast cells (34). SMAD3 has also been implicated in EVT development. It was shown that depletion of SMAD3 but not SMAD2 suppresses EVT emergence from first-trimester human placental explants (35), indicating a specific role of SMAD3 during EVT development. Interestingly, our scRNA-seq analyses with first-trimester human placentae showed that all of these MMPs as well as PLAC8 and SMAD3 are induced in developing EVTs. Thus, our findings indicate that WWTR-1 might mediate multi-pronged roles during EVT development by regulating expressions of MMPs, PLAC8 and SMAD3. Given the dynamic nature of EVT development and the essential role of EVTs in human placentation, it is important to institute future studies to better understand the role of WWTR1 in EVT development and function.

The loss of WWTR1 expression in CTBs from pregnancies with extreme preterm-birth and IUGR/PE indicates a direct correlation of WWTR1 with adverse pregnancies. We also showed that loss of WWTR1 in human TSCs and primary CTBs promotes a premature differentiation to STB lineage. Inductions of STB-specific gene expression were also noticed when TEAD4 and YAP1 were depleted in human TSCs and CTBs, respectively. Intriguingly, elevated maternal serum levels of human chorionic gonadotropin (hCG) and Inhibin-A, measured at 15–20 weeks gestation, increase the subsequent risk of IUGR/PE and extreme preterm birth (37). Since both hCG and Inhibin-A are produced by STBs and extreme preterm birth are often associated with loss of proliferating CTBs (37), it was proposed that elevated levels of hCG and/or Inhibin-A may result from premature differentiation of the CTBs to adopt STB fate (35). Thus, our findings from this study and prior studies with TEAD4 and YAP1 (11, 28) support the hypothesis that during human placenta development, loss of hippo signaling components, such as WWTR1, YAP1 and TEAD4, may result in premature accelerated differentiation of CTBs to STBs, which subsequently contributes to adverse pregnancies, like extreme preterm birth and IUGR/PE.

## EXPERIMENTAL PROCEDURES

### Human placental sample analysis

De-identified and discarded first-trimester placental tissues and term placental samples from normal and pathological pregnancies were obtained from Mount-Sinai Hospital, Toronto or collected at the University of Kansas Medical Center. The IRB at Mount Sinai Hospital and the University of Kansas IRB approved all collections and studies. Fresh first-trimester placental tissues were embedded in OCT and cryo-sectioned or used for scRNA-seq analyses. To test EVT development, first-trimester placental explants were cultured on matrigel for 6-8 days in medium that promote EVT differentiation in human TSCs (see below).

### Single-Cell RNA sequencing and analysis

Details of single-cell RNA-seq analyses with first-trimester placenta was performed and reported earlier (11). Briefly, single-cell suspensions from two first-trimester placentae were generated and transcriptomic profiles were obtained using the 10x Genomics Chromium Single Cell Gene Expression Solution (https://10xgenomics.com). The primary analysis of the scRNAseq data was performed using the 10x Genomics Cell Ranger pipeline (version 3.1.0). This pipeline performs sample de-multiplexing, barcode processing, and single cell 3’ gene counting. The quality of the sequenced data was assessed using the FastQC software. Sequenced reads were mapped to the human reference genome (GRCh38) using the STAR software. Individual samples were aggregated using the “cellranger aggr” tool in Cell Ranger to produce a single feature-barcode matrix containing all the sample data. The Cell Ranger software was used to perform t-SNE projections of cells, and k-means clustering. The 10x Genomics Loupe Cell Browser software was used to find significant genes, cell types, and substructure within the single-cell data. The raw data for scRNAseq analyses are submitted to the GEO database (http://www.ncbi.nlm.nih.gov/gds), with accession number GSE145036.

### Human TSC culture

Human TSC lines, derived from first trimester CTBs, were described earlier (11, 25). To maintain stem state culture, human TSCs were cultured on collagen IV-coated (5μg/ml) plate in DMEM/F12 medium, supplemented with 0.1 mM 2-mercaptoethanol, 0.2% FBS, 0.5% Penicillin-Streptomycin, 0.3% BSA, 1% ITS-X supplement, 1.5 μg/ml L-ascorbic acid, 50 ng/ml EGF, 2 μM CHIR99021, 0.5 μM A83-01, 1 μM SB431542, 0.8 mM Valproic acid and 5 μM Y27632. For EVT differentiation, TSCs were resuspended in 2% Matrigel (Corning, NY) and media containing DMEM/F12 supplemented with 0.3% BSA, 1% ITS-X, 0.5% Penicillin-Streptomycin, 100μM β-Mercaptoethanol, 2.5μM Y27632, 7.5μM A83-01, 100ng/ml hNRG1 and 4% Knockout serum. EVT differentiation medium without hNRG1 was replaced on day 3. On day 6 the media, lacking hNRG1 and KSR, was again replaced and finally analysed on day 8.

### RNA Interference (RNAi) in Human TSCs

Lentiviral shRNAs were used to knockdown *WWTR1* (target sequence: GCGATGAATCAGCCTCTGAAT) in human TSCs. A scramble shRNA (Addgene-1864, CCTAAGGTTAAGTCGCCCTCGC) was used as control. Lentiviral particles were generated by transfecting plasmids into HEK-293T cells. Virus containing supernatant was collected and virus particles were concentrated by Lenti-X concentrator (Clontech Laboratories, CA) according to the manufacturer protocol. Human TSCs were transduced using viral particles at 60-70% confluency. Transduced cells were selected in the presence of puromycin (1.5-2μg/ml). Selected cells were tested for knockdown efficiency and used for further experimental analyses.

### CTB Isolation from first trimester placenta

CTBs were isolated from 8-10^th^ week first-trimester pooled placentae (n=8) as described (28). Briefly placentas were kept overnight in DMEM HAM’s F12 (Gibco 31331-28)/ 0.05mg/mL gentamicin (Gibco 15710-049), 0.5μg/mL fungizone (Gibco 15290026). Next day placental villi were scraped in 1x Hank’s Balanced Salt Solution (HBSS, Sigma H4641) collected by centrifugation and incubated for two consecutive digestions with 1x HBSS containing 0.125% trypsin (Gibco 15090-046) and 0.125mg/mL DNase I (Sigma-Aldrich DN25) at 37°C in the incubator. Cells were purified by Percoll (cytiva 17089101) gradient centrifugation. Contaminating erythrocytes were lysed by incubation with erythrocyte lysis buffer (155mM NH_4_Cl, 10mM KHCO_3_, 0.1mM EDTA, pH 7.3) for 5min at room temperature. The cell suspension was seeded onto cell culture dishes for 45 min to allow contaminating stromal cells to adhere to the plastic. Trophoblasts were collected from the supernatants by centrifugation. HLAG^+^ EVTs were depleted from the cell suspension by immune-purification using HLA-G PE labeled antibodies (Exbio Clone MEM-G/9, 1P-292-C100), PE MACS beads (Miltenyi biotec 130-048-801) and MACS MS columns (Miltenyi biotec 130-042-201). Purified Trophoblasts were seeded in DMEM-HAM’s F12 (Gibco 31331-28)/ 10% FBS (Sigma S0615-500ML) 0.05mg/mL gentamicin (Gibco 15710-049), 0.5μg/mL fungizone (Gibco 15290026) onto fibronectin coated cell culture dishes (2μg/cm^2^, Merck FC010). For siRNA mediated gene silencing, one hour later a proportion of the cell culture media was replaced by siRNA/RNAimax containing media prepared as described (43) by using non targeting (D-001810-10-20) or TAZ (L-016083-00-0005) ON-TARGETplus SMARTpools and Lipofectamine RNAimax (Invitrogen 13778-075).

### Placental explants culture

First-trimester placental explants were submerged in DMEM/F12(Gibco) media and divided into smaller pieces under the dissection microscope in sterile conditions. Pieces containing branching villous like structure were washed in phosphate buffer saline (PBS) supplemented with 10% fetal bovine serum (FBS) and then subjected to lentiviral treatment. The explants were divided into two groups, one group was incubated with scrambled lentiviral particles while the other group was incubated with lentiviral particles carrying shRNA for WWTR1 gene knockdown. Both the groups were incubated with the respective lentiviral particles for 6hrs, at 37 °C in a humidified chamber in a 5% CO2/95% air gas mixture. After 6 hours the explant pieces were rinsed and encapsulated in gel for further culture. For the encapsulation, Growth factor reduced Matrigel (Corning) was mixed 1:1 with DMEM/F12 on ice to make matrigel suspension. 200ul of the matrigel suspension was added to each well of a 24-well plate and explants were placed centrally and covered with another 200ul of matrigel suspension. The plate was then incubated at 37 °C in a humidified chamber in a 5% CO2 for the gel-suspension to solidify thereby encapsulating the explant. Finally, 300ul of EVT media was added to each well and allowed to culture. EVT media was changed on days 3 and 5 as mentioned earlier.

### RNA-Seq analysis

RNA sequence analysis was performed according to published protocol (44, 45). Total RNA from the control human TSCs as well as *WWTR1-*KD human TSCs were isolated using RNeasy Mini Kit (Qiagen, 74104) per the manufacturer’s protocol with on column DNase digestion. RNA concentrations were quantified using a NanoDrop Spectrophotometer at a wavelength of 260nm. Integrity of the total RNA samples was evaluated using an Agilent 2100 Bioanalyzer (Agilent Technologies Inc., Santa Clara, CA). The total RNA fraction was processed by oligo dT bead capture of mRNA, fragmentation and reverse transcription into cDNA. After ligation with the appropriate Unique Dual Index (UDI) adaptors, cDNA library was prepared using the Universal Plus mRNA-seq +UDI library preparation kit (NuGEN 0508-08, 0508-32). The raw data for RNA-seq analyses are available at GEO database with accession number GSE188738.

### Statistical significance

Statistical significances were determined for quantitative RT-PCR analyses for mRNA expression and for cell proliferation analyses. We have performed at least n=3 experimental replicates for all of those experiments. For statistical significance of generated data, statistical comparisons between two means were determined with Student’s t-test and significantly altered values (p≤0.01) are highlighted in figures by an asterix. RNA-Seq data were generated with n=4 experimental replicates per group. The statistical significance of altered gene expression (absolute fold change ≥ 2.0 and FDR q-value ≤ 0.05) was initially confirmed with right tailed Fisher’s exact test. For *WWTR1* mRNA expression analyses in pathological placentae, one-way analysis of variance is used to determine statistically significant differences in mean WWTR1 expression in a specific pathological condition, such as extreme preterm birth, with term control placentae.

## Data Availability

The raw data for RNA-seq analyses are available at GEO database (http://www.ncbi.nlm.nih.gov/gds) with accession number GSE188738. The raw data for single-cell RNA-seq in first-trimester human placenta is also available in GEO database (accession number GSE145036).

## ACKNOWLEDGEMENTS

This research was supported by NIH grants **HD101319**, HD062546, HD0098880, **HD103161** and **HD102188** and a pilot grant under NIH Center of Biomedical Research Program (COBRE, P30GM122731) to Soumen Paul. We acknowledge the Genomics Core, the Imaging and Histology Core and the Bioinformatics Core of the University of Kansas Medical Center. We thank Drs. Hiroaki Okae and Takahiro Arima of Tohoku University Graduate School of Medicine, Japan, for sharing human TSC lines. We thank Ms. Brandi Miller for critical comments on the manuscript.

