## Supplemental Methods and Figures for "Hippo Signaling Cofactor, WWTR1, at the Crossroads of Human Trophoblast Progenitor Self-Renewal and Differentiation"

**Supplemental Information Inventory:**

| Items | Related to | General Description |
|---|---|---|
| **Supplementary Experimental Procedures with References** | | Describes experimental procedures and lists oligonucleotides and antibodies that are used for the study. |
| **Supplementary Figure Legends** | | |
| **Figure S1** | **Related to Figure 1** | |
| **Figure S2** | **Related to Figure 1** | |
| **Figure S3** | **Related to Figure 1** | |
| **Figure S4** | **Related to Figures 3,4,5** | |
| **Figure S5** | **Related to Figure 4** | |
| **Figure S6** | **Related to Figure 6** | |
| **Figure S7** | **Related to Figure 7** | |
| **Dataset S1** | **Related to Figure 3,4,5,6** | Gene Expression changes in *WWTR1*-KD human TSCs. |

### SUPPLEMENTARY EXPERIMENTAL PROCEDURES

**Bioinformatics Analyses of RNA-Seq data:** Quality assessment has been done using FastQC v0.11.9 (https://www.bioinformatics.babraham.ac.uk/projects/fastqc/). The reads were aligned using the subread-2.0.3 (1), after that reads count file were obtained using featureCounts (2) of subread-2.0.3. After that differential expression was obtained using DESeq2 (3). The mean insert length and standard deviation calculated using picard 2.26.6 (https://broadinstitute.github.io/picard/). All the analysis has been done using UCSC hg38 from iGenomes illumine as reference data. The heatmap was generated using screening top 100 genes by using "genefilter" library followed by pheatmap v1.0.12. MA-plot were generated using ggmaplot function of ggpubr v0.4.0 (https://cran.r-project.org/web/packages/ggpubr/index.html) using the parameters fdr = 0.05, fc = 2. The volcano plot was done using ggplot2 v3.3.5 (https://ggplot2.tidyverse.org/) where intercept were provided at log2fold values at (-1,1) to get the better view. GSEA analysis was performed using GSEA_Linux_4.2.2 (4) after preparing the two files with extension '.gct' and '.cls' generated as per prescribed guideline of tool. Gene sets c5.go.mf.v7.5.1.symbols.gmt was selected for analyses for GO molecular Function, while c2.cp.v7.5.1.symbols.gmt was selected for analysis for Canonical Pathways.

**Organoid generation from human trophoblast stem cells and cultivation:** Human TSCs and first-trimester CTBs were used to generate the self-renewing organoids. For CTB organoid, we followed earlier described protocols (5, 6). To obtain organoids from human TSCs (control and *WWTR1*-KD human TSCs) we modified the protocol described previously (5). Human TSCs were harvested and re-suspended in ice-cold basic trophoblast organoid medium (b-TOM) containing advanced DMEM/F12 supplemented with 10mM HEPES (Sigma H3537), B27 (Gibco 17504-044), N2 (Gibco 17502-048) and 2mM glutamine (Gibco 25030081). The cells were then centrifuged for 3 minutes at 1200rpm following which the cells were re-suspended in ice-cold advanced trophoblast organoid medium (aTOM) which is b-TOM supplemented with 100ng/ml R-spondin (PeproTech 120-38-20UG), 1µM A83-01 (Sigma SML0788), 100ng/ml recombinant human epidermal growth factor (rhEGF, Sigma E9644), 50ng/ml recombinant murine hepatocyte growth factor (rmHGF, PeproTech 315-23-20UG), 2.5µM prostaglandin E2 (R&D System 2296/10), 3µM CHIR99021 (Sigma SML1046) and 100ng/ml Noggin (Invitrogen PHC 1506). Growth factor reduced matrigel (Corning) was added to the a-TOM cell suspension to reach a final concentration of 60%. 35µl of the viscous cell solution containing $2.5 \times 10^4$ cells was plated in the center of a 24-well plate. The solution rests as a dome-shaped droplet in the center of the well. The plates are then turned upside down and kept at 37°C for 10-15 minutes to ensure proper spreading of the cells in the solidifying matrigel domes. Finally, the plates are returned to their upward position and the domes are overlaid with 500µl of room temperature a-TOM medium. The organoids are allowed to form for 8-10 days with fresh media being changed every 2 days. Brightfield images were taken to observe the growth of the organoids.

**Cell Proliferation Assay:** Proliferations of control and *WWTR1*-KD human TSCs were determined via two different approaches. In the first approac**h,** $2x10^5$ number of cells were seeded on a 6 well plate and cell counts were taken for control and WWTR1-KD human TSCs at 24, 48, and 72 hours. Cells were imaged and compared using a brightfield microscope (Olympus IX71, Japan). In the second approach, 1.5x104 number of cells were seeded on a 12 well plate and the cell proliferation assay was performed by using the 5-Bromo-2′ deoxy-uridine labeling and detection kit I (Roche 12296736001) (according to the manufacturer' s manual). The cells were cultured for 24, 48, 72 hours. Hoechst 33342 (5 µg/ml) prepared in slow fade (Invitrogen) was used to counter-stain the nuclei. The BrdU-positive cells were observed and calculated using a fluorescent microscope (Nicon Eclipse 80i).

**Immunofluorescence (IF) and Immunohistochemistry (IHC) study:** Immunofluorescence was performed using human placenta cryosections. The sections were fixed using 4% paraformaldehyde in 1X PBS (sigma D8537), permeabilized using 0.25% Triton X-100 (Sigma X100) in 1X PBS and blocked for 1 hour using blocking buffer (10% FBS and 0.1% TritonX100 in PBS). Sections were incubated with primary antibodies overnight at 4 °C, washed with 0.1% Triton X-100 in PBS. After incubation (1:400, 1 h, room temperature) with conjugated secondary antibodies, sections were washed and mounted using antifade mounting medium (Thermo Fisher Scientific) containing DAPI. Immunohistochemistry was performed using paraffin sections of human placenta. The slides were deparaffinized by histoclear (National diagnostics, HS-200) and subsequently rehydrated with 100%, 90%, 80% and 70% ethanol. Antigen retrieval was done using Decloaking chamber at $80^0$C for 15 minutes. The endogenous peroxidase was inactivated by treating with 3% $H_2O_2$ (Sigma 216763) followed by wash with 1X PBS. Non-specific immunoglobulin binding was blocked with 10% goat serum (Thermo Fisher scientific-50062Z) for 1hour at RT followed by overnight incubation with 1:100 dilution of primary antibody or IgG at $4^0$C. The slides were washed 3 times with 1X PBS and incubated with biotin conjugated secondary antibody (1:200 dilution) for 1 hour at RT. The slides were washed with 1X PBS followed by treatment with streptavidin conjugated horseradish peroxidase (Vector laboratories, CA, SA-5704) for 20 minutes at RT. Reactivity was detected using DAB+ substrate chromogen system (Dako, TA-125-QHDX). The slides were counterstained with Mayer's hematoxylin (Sigma MH516). The slides were then dehydrated by sequential treatment using 70%, 80%, 90%, 100% ethanol and xylene. The sections were dried, mounted with Toluene and imaged using Nikon TE2000 microscope. Antibodies used for staining are as follows; (i) CDH1 (HECD-1), ab1416 (Abcam), (ii) Wnt7a, ab100792 (Abcam), (iii) WWTR1, HPA007415 (Sigma) (iv) P40 (TP63), ACI3066A (BioCare Medical, CA) and (v) CGB (hCGβ ), ab53087 (Abcam).

**Western blot analyses:** Protein lysates and supernatants of cell cultures were separated on SDS/PAA gels, transferred to Hybond-P PVDF membranes (Amersham 10600023) and incubated with antibodies following an earlier described protocol (7). Antibodies used for the study are as follows; (i) WWTR1, 23306-1-AP (ProteinTech, IL), (iii) Wnt7a, ab100792 (Abcam, Cambridge, MA), (iv) β-Actin, A5441 (Sigma, St.Louis, MO), (v) GAPDH: Cell Signaling 2118, (4) CGB: Dako A0231). HRP-conjugated secondary antibodies, the WesternBright Chemilumineszenz Substrat Quantum (Biozym 541015) and a ChemiDoc Imaging System (Bio-Rad) were used for signal visualization.

**Chromatin Immuno-precipitation (ChIP) assay:** Quantitative ChIP analyses were performed to determine WWTR1 occupancy at selected gene loci in Human TSCs. We followed a published protocol (8). Cells were crosslinked with 1% formaldehyde (Sigma) for 10 mins at room temperature with gentle rotation. Chromatin crosslinking was stopped with glycine (125mM). These samples were sonicated. Chromatin fragments were immunoprecipitated with 8 microgram of WWTR1 antibody (HPA007415 Sigma) per ChIP experiment.

Primers used for Real-Time PCR analysis

| Primer Name | Forward | Reverse |
|---|---|---|
| TP63 | GTCATTTGATTCGAGTAGAGG GG | CTGGGGTGGCTCATAAGG T |
| CGB | GTGTGCATCACCGTCAACAC | GGTAGTTGCACACCACCTGA |
| CGA | TCTGGTCACATTGTCGGTGT | TTCCTGTAGCGTGCATTCTG |
| PSG4 | CGATGGGACTGGAGGAGTAA | AGTTGCTGCTGGAGATGGAG |
| ERVFRD-1 | CCAAATTCCCTCCTCTCCTC | CGGGTGTTAGTTTGCTTGGT |
| MMP2 | TCTCCTGACATTGACCTTGGC | CAGGGTGCTGGCTGAGTAGATC |
| Wnt7a | CTGTGGCTGCGACAAAGAGAA | GCCGTGGCACTTACATTCC |
| Wnt3 | AGGGCACCTCCACCATTTG | GACACTAACACGCCGAAGTCA |

| | | |
|---|---|---|
| Wnt4 | GTACGCCATCTCTTCGGCAG | GCGATGTTGTCAGAGCATCCT |
| WWTR1 | TCCCAGCCAAATCTCGTGATG | AGCGCATTGGGCATACTCAT |
| HPRT1 | ACCCTTTCCAAATCCTCAGC | GTTATGGCGACCCGCAG |
| MMP2 | TCTCCTGACATTGACCTTGGC | CAGGGTGCTGGCTGAGTAGATC |
| MMP11 | CCGCAACCGACAGAAGAGG | ATCGTCCCATACCTTTAGGGC |
| MMP14 | CGAGGTGCCCTATGCCTA | CTCGGCAGAGTCAAAGTGG |
| MMP15 | AGGTCCATGCCGAGAACTG | GTCTCTTCGTCGAGCACACC |
| WWTR1 | TCCCAGCCAAATCTCGTGATG | AGCGCATTGGGCATACTCAT |
| HPRT1 | ACCCTTTCCAAATCCTCAGC | GTTATGGCGACCCGCAG |
| PLAC8 | TGCTAGAAAAAGCCATGGAA | TCTGCCAGAGGGTCTTTAGG |
| YAP1 | TCCACCAGTGCAGCAGAATA | TTCCCATCCATCAGGAAGAG |
| SMAD3 | TGAGGCTTATTAAGTCCATTGC | TCATCATTTGTCATACTGCACAC |

**References mentioned in the Supplementary Methods:**

**Fig. S1**

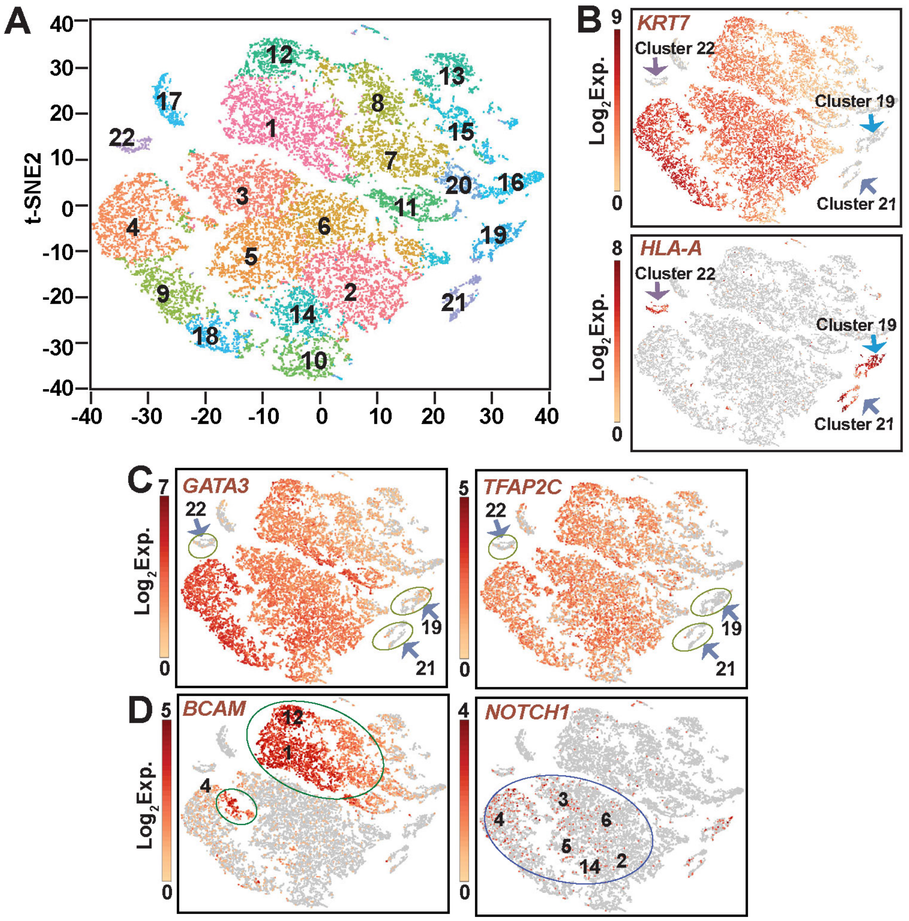


**Fig. S1. scRNA-seq analyses in first-trimester human placentae.** (A) The t-SNE plot of the aggregate of the hierarchical clustering of 2 first-trimester human placental samples (one Week 7 and one Week 8) identified 22 different cell clusters. (B) Clusters of trophoblast and non-trophoblast cells are labeled by *KRT7* and *HLA-A* expressions, respectively, on a t-SNE plot. Lack of KRT7 expression and induction of *HLA-A* expression identified clusters 19, 21 and 22 as non-trophoblast cells. (C) t-SNE plots show almost all cells within trophoblast cell clusters express *GATA3* and *TFAP2C*, which are known to be expressed in all mononuclear trophoblast cell within a first-trimester human placenta. (D) t-SNE plots show expression patterns of *BCAM* and *NOTCH1*, which are predominantly expressed in stem-state CTBs and proximal column CTBs, respectively.

### Fig. S2

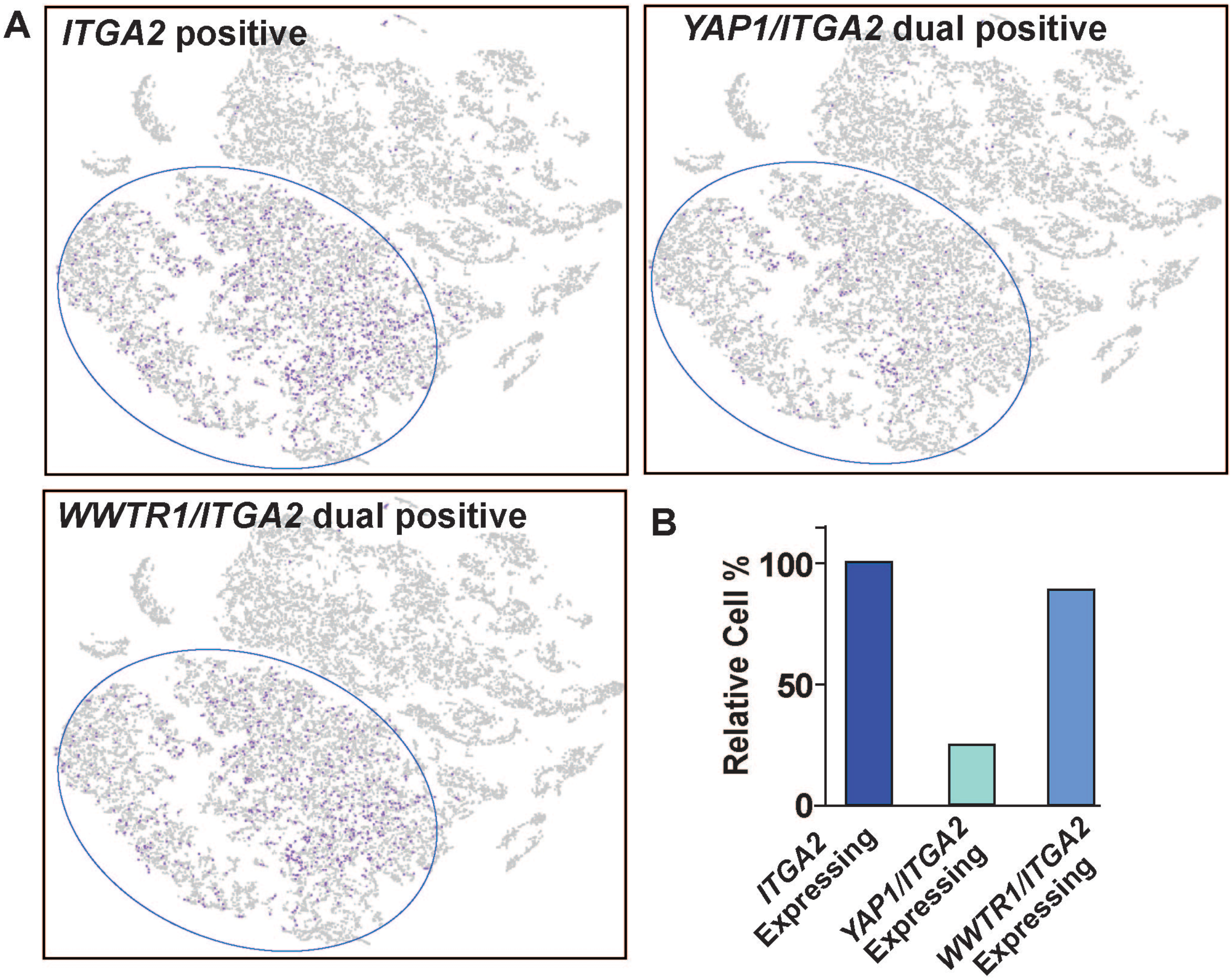


**Fig. S2. WWTR1 and YAP1 expression in ITGA2 positive CTB progenitors.** (A) and (B) *WWTR1* and *YAP1* expression was monitored and quantitated in *ITGA2*-expressing column CTB progenitors of a first-trimester human placenta via scRNA-seq analyses. Each blue dot represents CTB progenitors which are either *ITGA2* positive (A, upper left panel) and are also expressing *YAP1* (A, upper right panel) or *WWTR1* (A, Lower panel). Note that almost all *ITGA2*-expressing CTB progenitors also express *WWTR1*. However, only a fraction of *ITGA2* positive cells within a first-trimester placenta also express *YAP1*.

Fig. S3

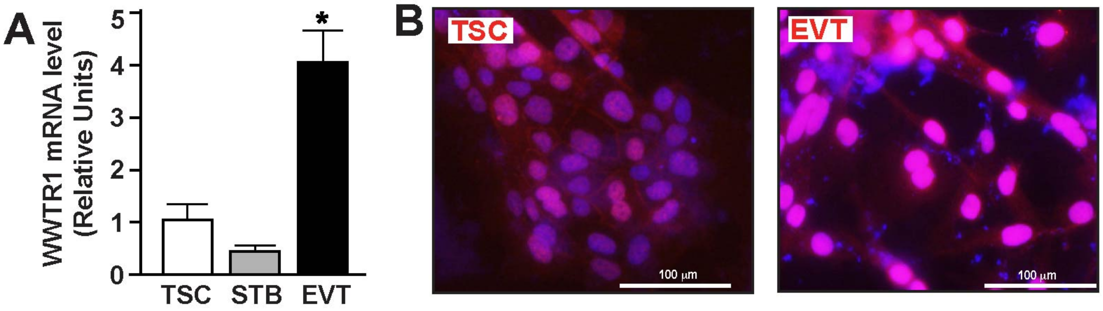


**Fig. S3. WWTR1 expression in human TSCs.** (A) RT-qPCR analyses showing relative expressions of *WWTR1* mRNA in undifferentiated human TSCs and after STB and EVT differentiation. (B) Immunofluorescence images show nuclear localization of WWTR1 in human TSCs when maintained at stem-state and after EVT differentiation.

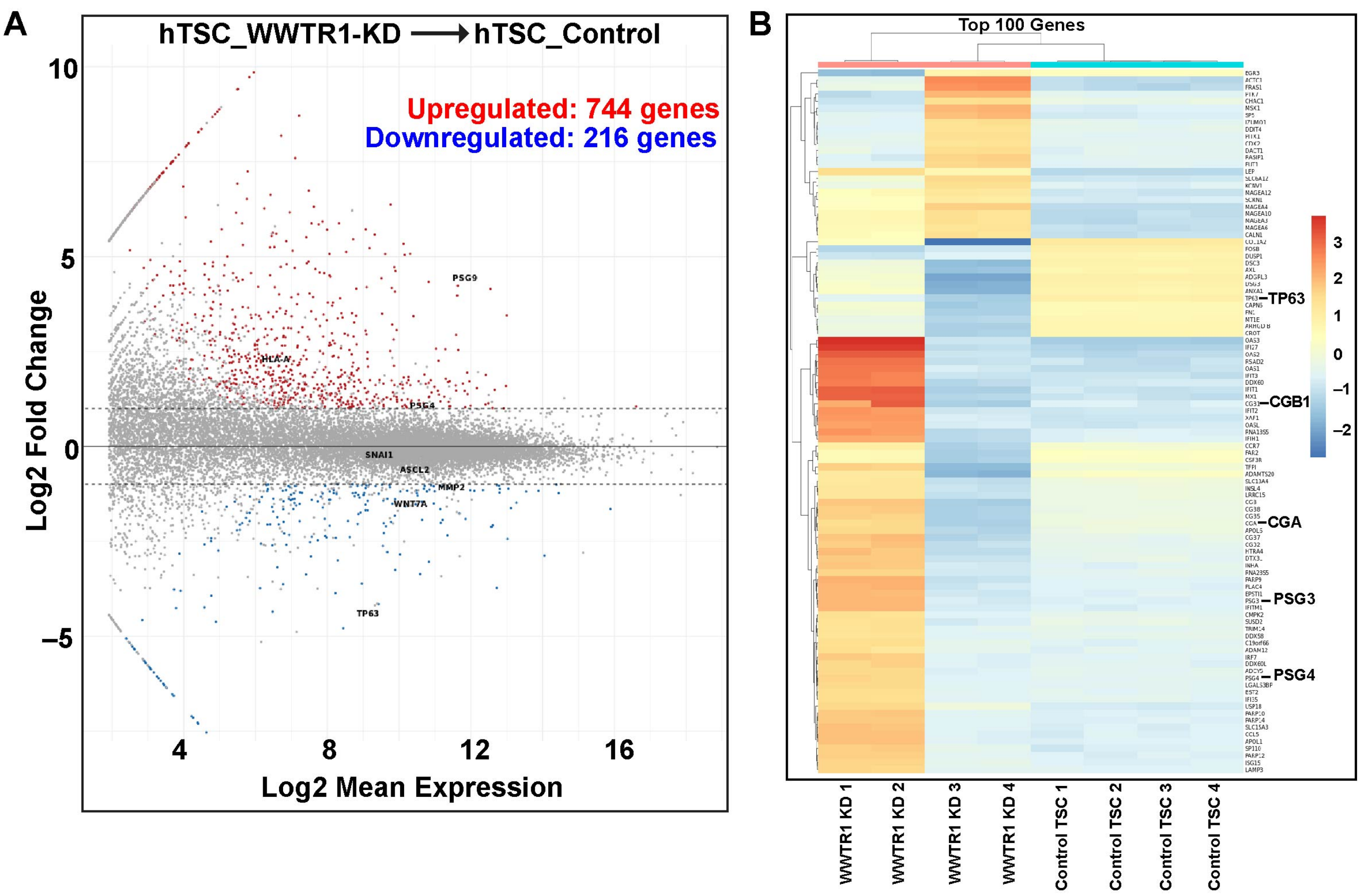


**Fig. S4. Global Genen Expression in WWTR1-KD human TSCs.** (A) Bland-Altman plot showing changes in global mRNA expression in human TSCs upon WWTR1 depletion. Colored dots indicate ≥2-fold change in mRNA expression with a false discovery rate of $P < 0.05$; red: up-regulated genes ( total 744 genes), blue: down-regulated genes (total 216 genes). (B) The heat map shows top 100 differentially expressed genes in *WWTR1*-KD human TSCs.

**Fig. S5**

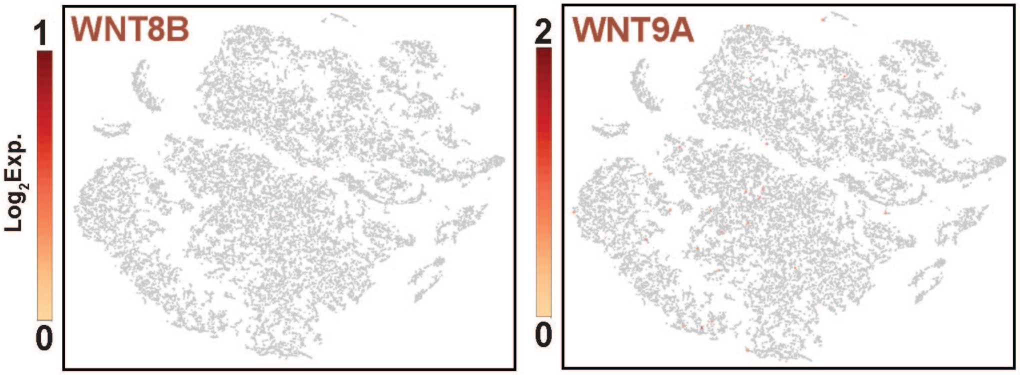


**Fig. S5.** t-SNE plots show lack of WNT8B and WNT9A mRNA expression in single cell clusters from first-trimester human placenta.

### Fig. S6

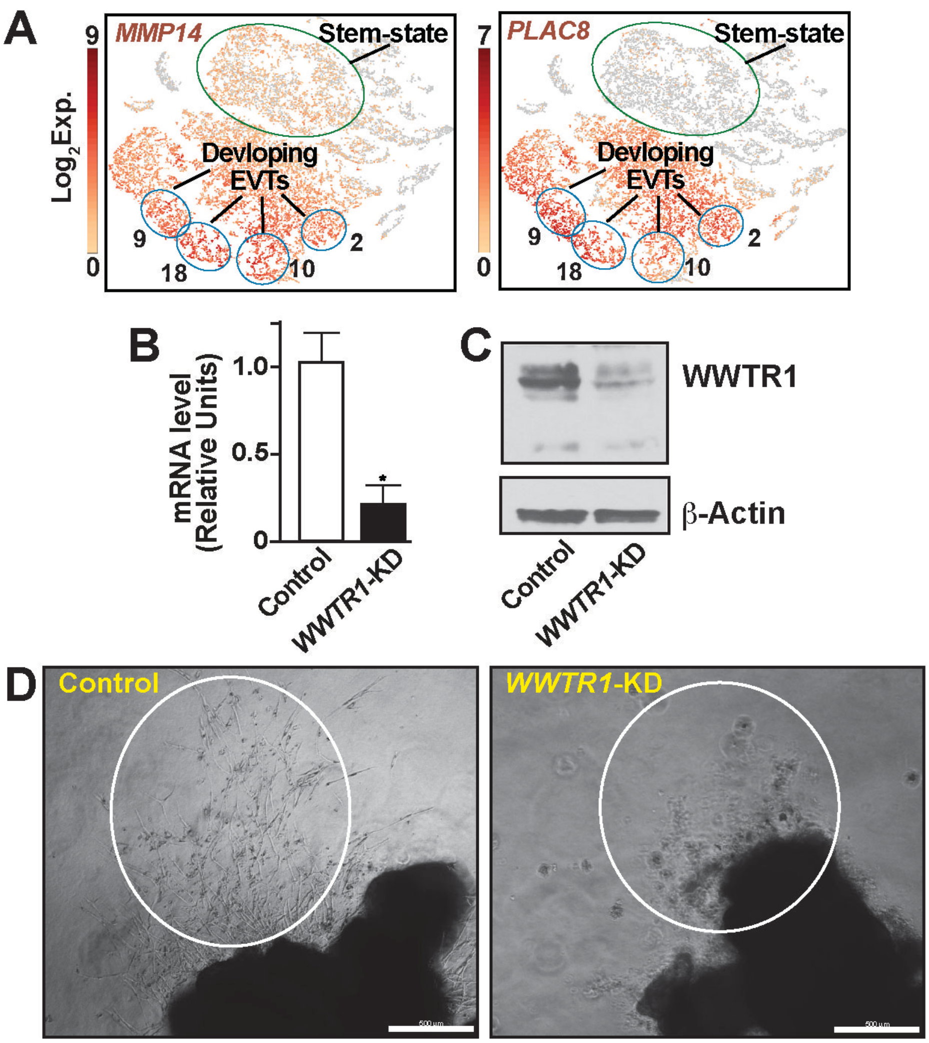


**Fig. S6.** (A) t-SNE plots showing mRNA induction of *MMP14*, and *PLAC8* in single cell clusters representing developing EVTs of first-trimester human placentae (B) and (C) RT-qPCR and Wester Blot analysis, respectively, showing depletion of WWTR1 expression in first-trimester human plaecntal explants via shRNA-mediated RNAi. (C) Phase contrast images showing inhibition of EVT development (white elipses) from first-trimester human placental explants after RNAi-mediated depletion of WWTR1.

**Fig. S7**

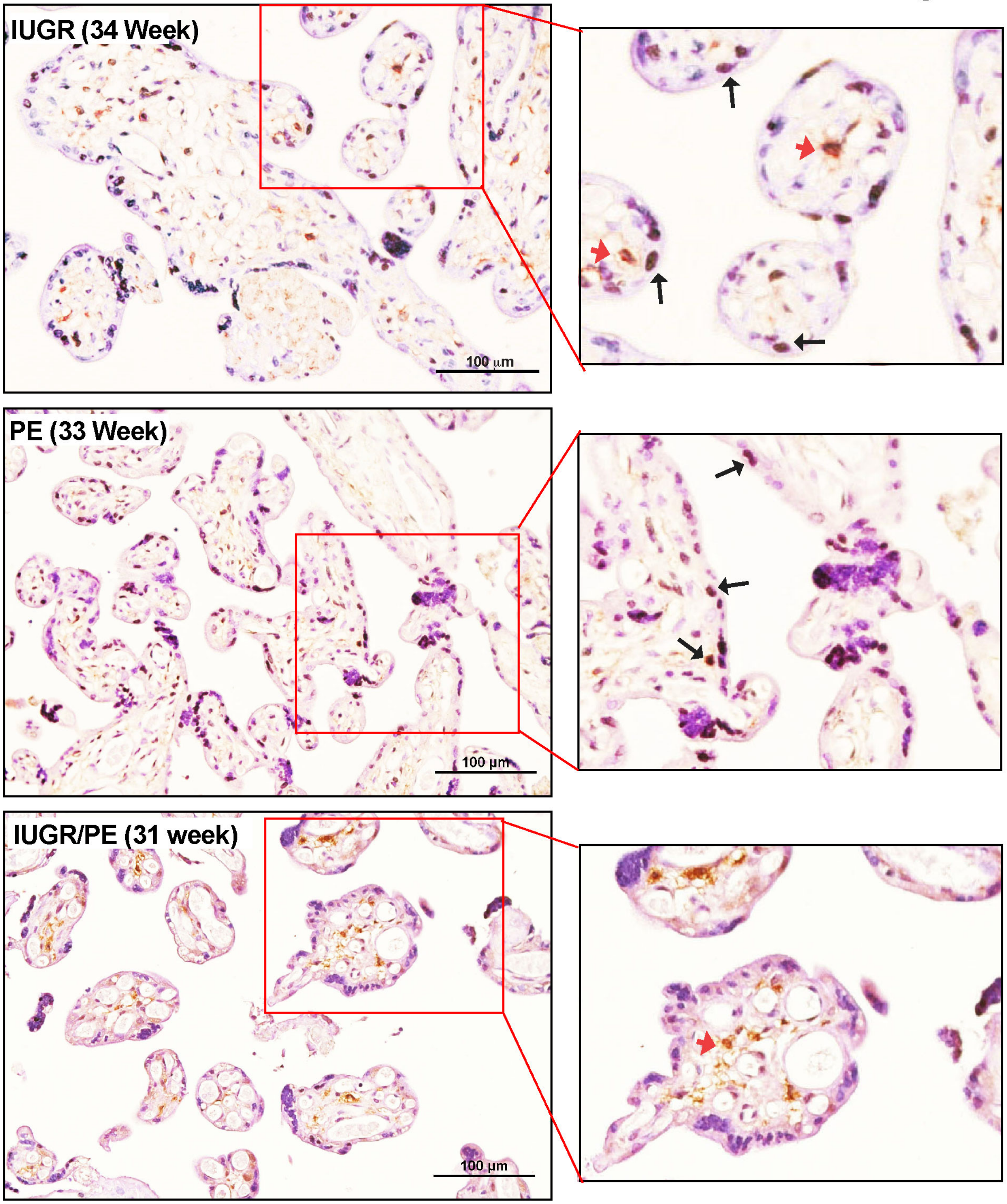


**Fig. S7. WWTR1 expression in pathological pregnancies** Representative immunostained images show WWTR1-expressing CTBs (Black arrows) in placentae from pregnancies that are associated with preterm birth in association with IUGR, PE or IUGR/PE. Red arrows indicate WWTR1-expressing non-trophoblast cells.
